# Substrate recognition by the *Pseudomonas aeruginosa* EF-Tu methyltransferase EftM

**DOI:** 10.1101/773556

**Authors:** Emily G. Kuiper, Debayan Dey, Paige A. LaMore, Joshua P. Owings, Samantha M. Prezioso, Joanna B. Goldberg, Graeme L. Conn

## Abstract

*Pseudomonas aeruginosa* is an opportunistic pathogen and a leading cause of serious infections in individuals with cystic fibrosis, compromised immune systems, and severe burns. During infection, *P. aeruginosa* adhesion to host epithelial cells is enhanced by surface exposed translation elongation factor EF-Tu carrying a Lys5 trimethylation. This modification is incorporated by the *S*-adenosyl-L-methionine-dependent methyltransferase EftM. Thus, EF-Tu modification by EftM may represent a novel target to restrict the establishment of *P. aeruginosa* infections in vulnerable individuals. Here, we extend our understanding of EftM action by defining the molecular mechanism of EF-Tu substrate recognition by this enzyme. First, following the observation that EftM can bind to EF-Tu lacking an N-terminal peptide (encompassing the Lys5 target site), an EftM homology model was generated and used in protein-protein docking studies to predict EftM:EF-Tu interactions. The predicted protein-protein interface was then experimentally validated using site-directed mutagenesis of residues in both proteins coupled with binding and methyltransferase activity assays. We also show that EftM is unable to methylate the isolated N-terminal EF-Tu peptide and that binding-induced conformational changes in EftM are likely needed to allow placement of the first 5-6 amino acids of EF-Tu into the conserved peptide binding channel. In this channel, a group of residues that are highly conserved in EftM family proteins position the N-terminal sequence to facilitate modification of Lys5. Our findings provide detailed insights into substrate recognition by this lysine methyltransferase, paving the way for a deeper understanding of EftM’s mechanism of action on EF-Tu.

---

Protein post-translational modifications (PTMs) add an additional level of complexity to the proteome and are critical to the biological functions of proteins in all domains of life. In bacteria, PTMs such as methylation are used to respond to environmental cues and may be critical for adaptation to or evasion of host immune systems during infection (1-3).

The bacterial methyltransferase CheR serves as the paradigm for environmental adaptation behavior arising from protein methylation (4). In many enteric bacteria, response to chemical stimuli occurs via *O*-methylation of glutamic acid residues on chemotaxis receptors by CheR. Bacterial methyltransferases can also directly influence host gene expression and host-pathogen interaction. For example, *Mycobacterium tuberculosis* expresses a secreted 5-methylcytosine-specific DNA methyltransferase (Rv2966c) that directly modifies the host epigenetic machinery to alter transcription (5). Establishment of infection can also be both negatively and positively regulated by methylation of bacterial surface proteins by host- or pathogen-derived methyltransferases. The human protein SUV39H1, for example, methylates the crucial surface-exposed mycobacterial protein HupB which reduces bacterial adhesion and survival inside the host cell (6). In rickettsia, methylation of outer membrane protein B (OmpB) directly controls virulence (7): virulent species display lysine trimethylation of OmpB, which mediates host adhesion, attachment, and invasion, whereas avirulent strains possess monomethylated OmpB. Finally, our previous work has revealed a role in establishment of *P. aeruginosa* infection for the trimethylation of translation elongation factor thermo-unstable (EF-Tu) on its Lys5 residue by the EF-Tu methyltransferase, EftM (8-10).

While roles for protein methylation in bacterial virulence and host-pathogen interactions are emerging, there is generally less known regarding the molecular mechanisms that control substrate recognition and catalysis by the bacterial methyltransferase enzymes that incorporate these PTMs. The Gram-negative environmental bacterium *P. aeruginosa* is an opportunistic pathogen that causes serious respiratory, eye, and skin infections in vulnerable individuals, such as those with cystic fibrosis, compromised immune systems, or severe burns. *P. aeruginosa* is also intrinsically resistant to many antibiotics making infections difficult to treat once established. However, precisely how *P. aeruginosa* establishes infection in a host and adapts to its new environment is still not fully understood and a more complete understanding of this process could potentially identify novel strategies to restrict the establishment of infection. Toward this goal, we previously described trimethylation of EF-Tu on Lys5 as a PTM that appears to enhance *P. aeruginosa* host cell adherence and infection (8,9). In *P. aeruginosa* isolates from acute infections, as well as the laboratory strain PAO1, this EF-Tu modification is present in bacteria cultured at 25 °C but absent 37 °C (8,9,11). This thermoregulation of PTM incorporation was found to occur at the level of transcription initiation as well as the direct thermolability of the modifying enzyme, EftM (8,10). In contrast, some chronic isolates, such as PAHM4, retain their ability to methylate EF-Tu at 37 °C (11). However, whether this arises from increased thermal stability of EftM has not been tested. Together, these observations suggest that the major role of the Lys5 PTM may be to modulate a “moonlighting” function of EF-Tu in host cell adhesion by *P.aeruginosa*.

Here, we define the mechanism of EF-Tu substrate recognition by EftM. Evolutionary sequence analyses and computational modeling of EftM and the EftM:EF-Tu complex were used to direct mutagenesis and functional studies to identify the protein-protein binding interface, revealing that EftM requires more than just the N-terminal sequence of EF-Tu to bind and methylate its substrate. We also identify conserved residues in a substrate binding channel that position the N-terminal sequence of EF-Tu for trimethylation on Lys5. Collectively, our findings lead to a new mechanistic model for EftM:EF-Tu recognition reliant on both protein-protein docking and protein conformational changes to accommodate the EF-Tu N-terminal sequence, which together direct specific substrate recognition and modification. These functional studies represent an important step toward a better understanding of the molecular recognition mechanism of EftM and bacterial lysine methyltransferases in general.

## RESULTS

### EftM^HM4^ is functionally equivalent to EftM^PAO1^ but with increased thermostability

We previously reported that the temperature sensitivity of EF-Tu methylation by EftM in *P. aeruginosa* strain PAO1 (EftM^PAO1^) is due, at least in part, to the direct unfolding of the methyltransferase upon transition from environment-like (25 °C) to host-like (37 °C) temperature (8). This direct thermolability also likely contributed to challenges we encountered in expression, purification and handling of EftM^PAO1^ for biochemical and biophysical studies. For the current work, we anticipated such issues would likely be exacerbated for some EftM^PAO1^ variants made to investigate the enzyme’s interaction with EF-Tu. Therefore, we decided to examine the folding and functional properties of the same enzyme (EftM^HM4^) from the chronic infection strain of *P. aeruginosa* PAHM4 in which EF-Tu methylation was previously observed at 37 °C (9).

Using differential scanning fluorimetry (DSF) we found that the T_m_ of EftM^HM4^ is increased by ∼5° to 37 °C (**Fig. 1*A***), compared to 31.5 °C which we determined previously for EftM^PAO1^ (8). Additionally, as observed for EftM^PAO1^, *S*-adenosyl-L-methionine (SAM) stabilizes EftM^HM4^ by ∼5 °C resulting in an unfolding T_m_ of 41.5 °C in the presence of the cosubstrate. Next we asked whether this additional thermal stability of EftM^HM4^ directly correlates with its activity at elevated temperature. As expected, EftM^HM4^ retained enzymatic activity after incubation at either 25 °C and 37 °C as previously observed, but its activity is ablated after incubation at 42 °C for 5 minutes or more **(Fig. 1*B***). Collectively, these data show the retained methylation of EF-Tu in PAHM4 at 37 °C is due to an increase in the thermostability of EftM methyltransferase by ∼5°, which allows the enzyme to remain active at the elevated temperature. Critically, these findings also show that EftM^PAO1^ and EftM^HM4^ are functionally equivalent enzymes and that the latter still exhibits thermoregulation but shifted to higher temperature.

**Fig. 1.**
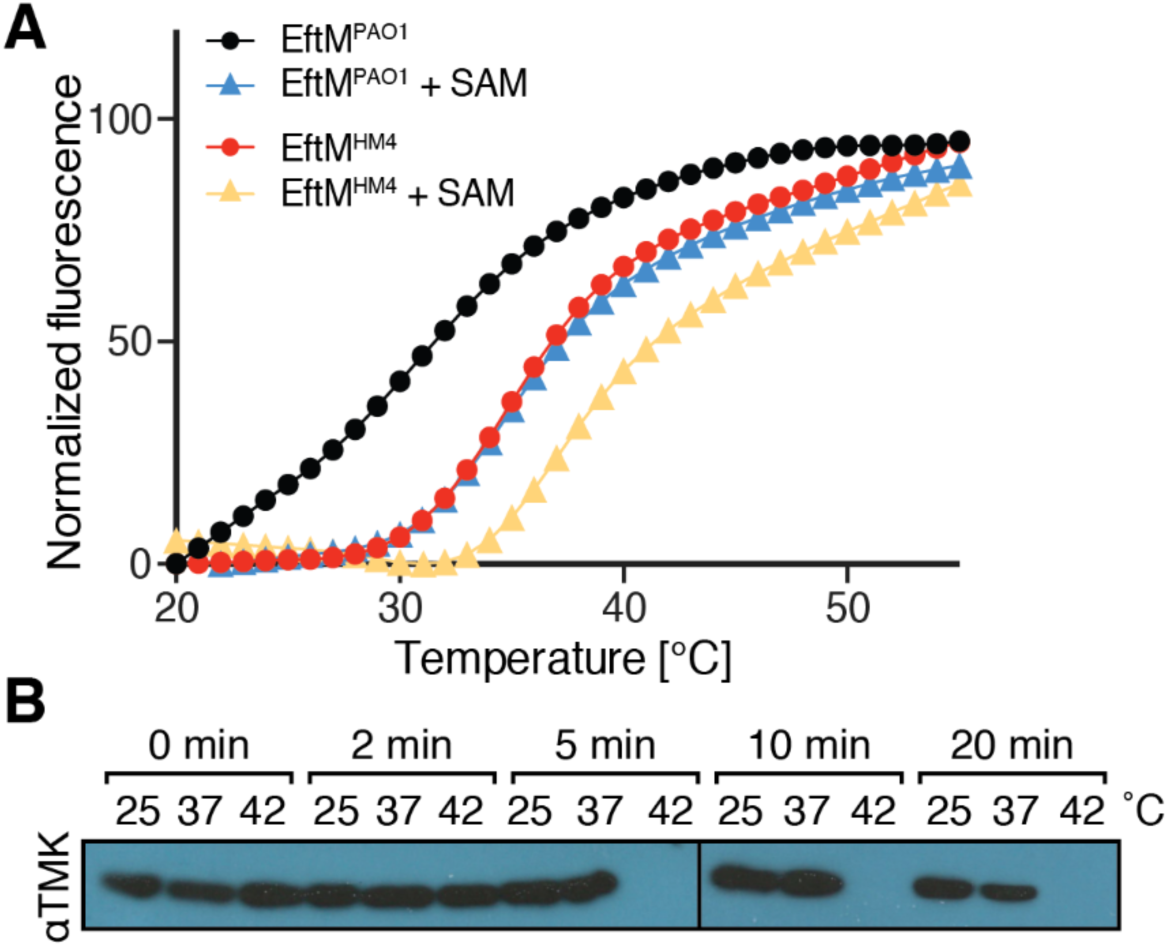
EftM from *P. aeruginosa* clinical isolate strain PAHM4 is more thermodynamically stable compared to EftM from PAO1 strain. ***A***, Differential scanning fluorimetry of EftM^HM4^ (red, circle) and EftM^PAO1^ (black, circle) alone or in the presence of 1 mM SAM (EftM^HM4^, tan triangle; EftM^PAO1^blue triangle). The data for EftM^PAO1^ were previously reported in (8) and shown here for comparison with EftM^HM4^. ***B***, *In vitro* methylation activity of EftM^HM4^ after incubation at a designated temperature.

As there is no known structure of any EftM family member, we searched for structures of the closest paralogs and identified the *N,N*-dimethyltransferase DesVI from *Streptomyces venezuelae* (PDB ID 3BXO), which methylates a sugar intermediate in macrolide antibiotic biosynthesis, dTDP-3-amino-3,6-dideoxy-*α*-D-glucopyranose (12). DesVI is 18.8% identical and 29.6% similar to EftM^HM4^, with both values ∼1% higher in comparison with EftM^PAO1^. A combined approach of homology modeling and threading, followed by energy minimization, was used to generate structural models of EftM^PAO1^ and EftM^HM4^. The essentially identical modeled EftM structures each consist of two domains: a Class I (Rossman fold) methyltransferase domain containing a central seven-stranded β-sheet and an auxiliary domain comprising the N-terminal *α*-helix and a three-stranded antiparallel β-sheet formed by an extended sequence that links the fifth and sixth strands (β5/6 linker) of the core methyltransferase fold **(Fig. 2**). The EftM models displayed the expected similarities with the structure of DesVI modeling template but also revealed some important potential differences. In particular, the substrate-binding channel of EftM is substantially enlarged, consistent with its binding of an extended substrate (*i.e.* the N-terminal sequence of EF-Tu). As the structure of DesVI was solved with the methyltransferase cosubstrate SAM, the model also revealed the likely location of cosubstrate and the surrounding catalytic center of EftM (**Fig. 2*B*,C**).

**Fig. 2.**
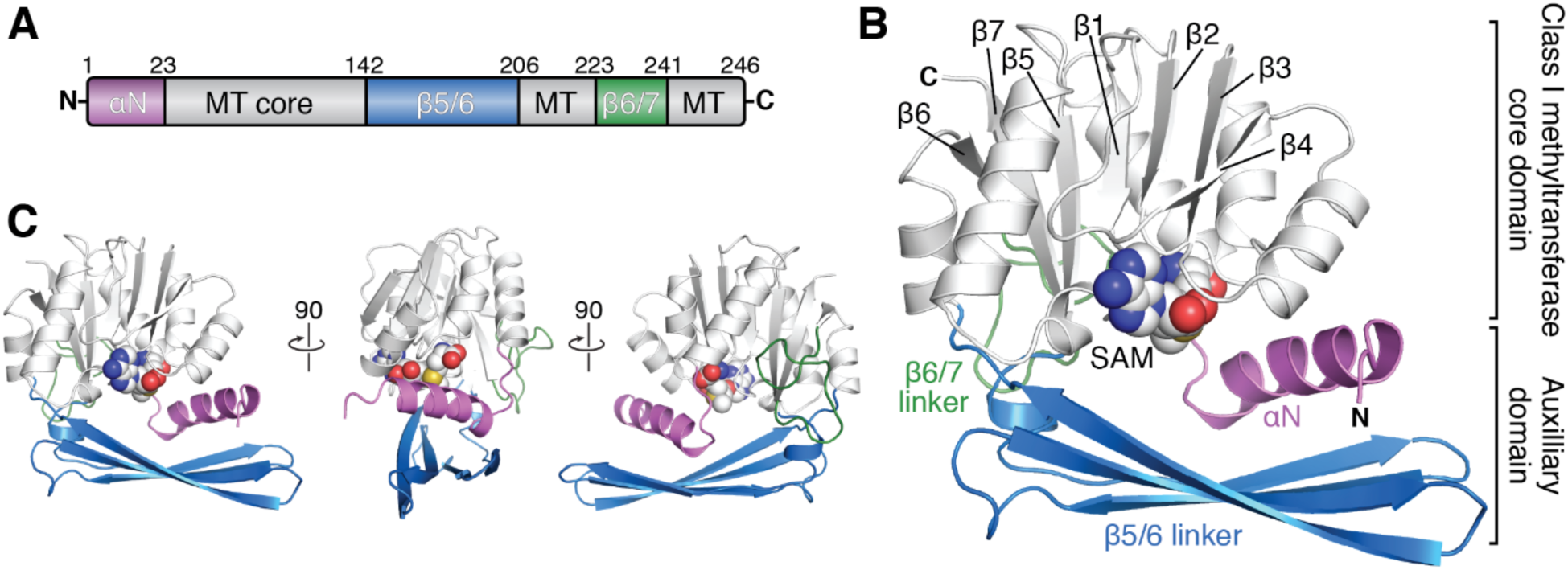
Homology model of EftM and structural elements. ***A***, Linear schematic of EftM structure highlighting the core Class I methyltransferase fold (MT; gray) and other regions, N-terminal extension (*α*N; pink), β5/6 linker (β5/6; blue), and β6/7 linker (β6/7; green). ***B***, Homology model of EftM^HM4^ modeled with SAM (from PDB 3BXO; shown as spheres) and its structural elements that embellish the conserved Class I methyltransferase core domain color coded as in *panel A*. ***C***, Three additional orthogonal views of the EftM homology model.

The amino acid sequences of EftM^HM4^ and EftM^PAO1^ are 96.4% similar and 88.7% identical, with 28 amino acids that vary between the two enzymes (**Fig. 3**). Almost all these 28 amino acid differences occur on the periphery of the enzyme suggesting that a network of interactions may be responsible for the increased stability of the EftM^HM4^, rather than one or a small number of individual amino acid changes. However, it is noteworthy that the changes involving the most physicochemically dissimilar amino acids are clustered in a region that connects the core methyltransferase fold and the auxiliary domain in the EftM model (**Fig. 3**, *red spheres*). This suggests a possible difference in regulation of protein flexibility between EftM^HM4^ and EftM^PAO1^, which might be responsible, at least in part, for the thermostabilization of EftM^HM4^.

**Fig. 3.**
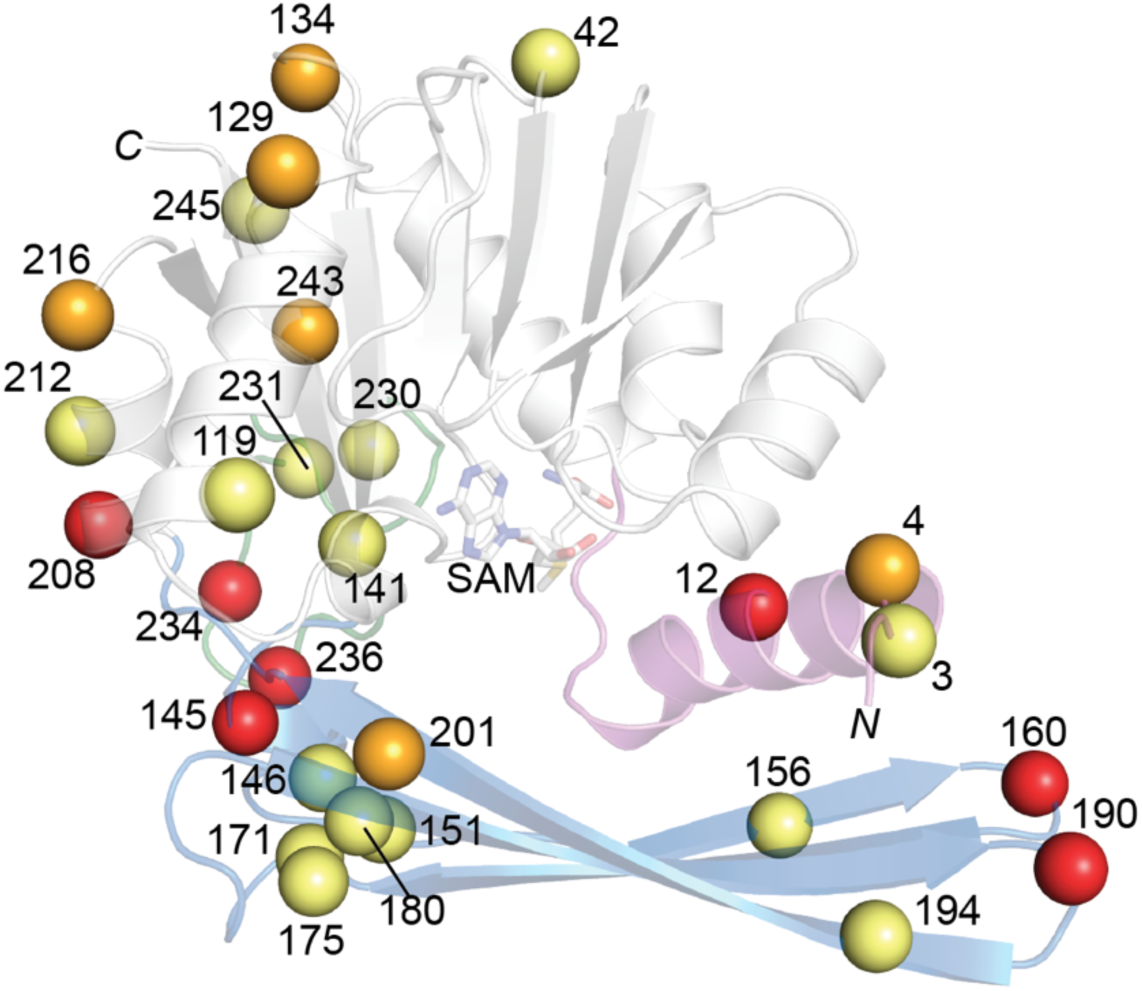
Differences between EftM^PAO1^ and EftM^HM4^ are distributed throughout the protein structure. Homology model of the EftM^HM4^-SAM complex with amino acid differences between EftM proteins from strains PAHM4 and PAO1 shown as spheres. Color coding indicates differences in amino acid physicochemical properties at each site: most distinct (red), intermediate (orange) and most similar (yellow).

In summary, EftM^HM4^ exhibits conserved activity, retained thermoregulation but with increased thermostability, and is predicted to have identical structure in comparison to the previously characterized thermolabile methyltransferase EftM^PAO1^. Coupled with its greater ease of expression, purification and handling, these factors lead us to use EftM^HM4^ as the optimal context to complete the detailed study described herein of EF-Tu recognition by EftM.

### EftM requires more than just the EF-Tu N-terminal peptide for binding

To begin defining EF-Tu recognition by EftM, we first asked whether important interactions are made with the surface of the globular protein fold of EF-Tu or if the isolated N-terminal peptide (^1^MAKE<u>**K**</u>FERNKP^11^; trimethylated Lys5 is underlined) alone is a suitable substrate for modification. First, a variant of *P. aeruginosa* EF-Tu lacking the first 11 amino acids of the N-terminus (Δ11-EF-Tu) was generated to assess contributions of the EF-Tu surface to binding by EftM. To qualitatively assess binding, we compared the ability of EftM to form a stable complex with EF-Tu or Δ11-EF-Tu using gel filtration chromatography (**Fig. 4*A,B***). Complexes of both EftM:EF-Tu and EftM:Δ11-EF-Tu were observed to elute from the column before either free component due to their larger size, suggesting that EftM is able to bind the truncated EF-Tu protein.

**Fig. 4.**
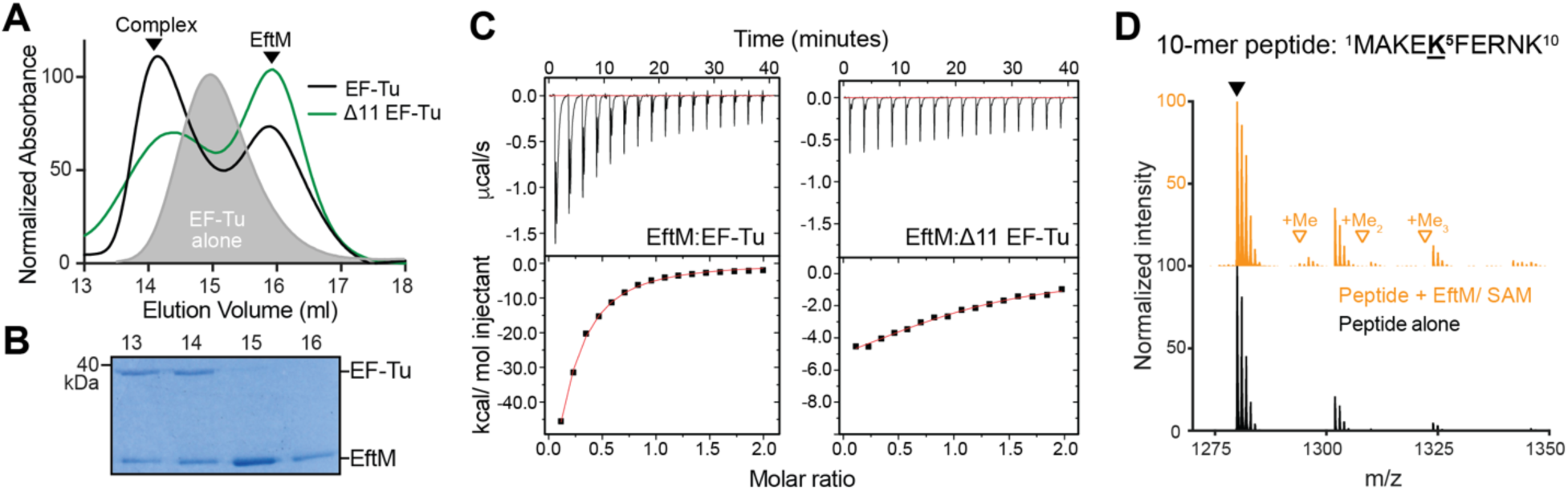
EftM recognizes and binds to more than just the EF-Tu amino terminus. ***A***, Gel filtration chromatography of EftM:EF-Tu complexes: EftM:EF-Tu (black), EF-Tu alone (grey shaded), EftM:Δ11-EF-Tu (green). ***B***, Coomassie-stained SDS-PAGE gel of gel filtration fractions confirming elution of the EftM:EF-Tu in the first peak in the chromatogram (fractions at 13 and 14 ml). ***C***, ITC analysis of binding of EftM and EF-Tu /Δ11-EF-Tu. Although weak binding affinity precludes accurate determination of *Δ*H, *i.e.* the intercept of the fit curve on the y-axis in the lower plot for each titration, note the difference in the scale of this plot for each complex which suggests that enthalpy of binding is significantly reduced in the absence of the EF-Tu N-terminal peptide in Δ11-EF-Tu. ***D***, MS analysis of a 10-mer peptide corresponding to the N-terminal peptide without (black) or with (orange) treatment with EftM and SAM in a methylation reaction.

We next quantified the binding affinity of EftM to wild-type *P. aeruginosa* EF-Tu and the Δ11-EF-Tu truncated construct using isothermal titration calorimetry (ITC). The measured binding affinity (K_D_) was relatively weak (∼27 μM) but found to be equivalent for both EF-Tu proteins (**Table 1**), consistent with retained interaction even in the absence of the EF-Tu N-terminal sequence. Interestingly, however, while the weak binding precludes accurate determination of binding enthalpy (ΔH), binding of EftM to Δ11-EF-Tu appears to have a significantly lower ΔH (>10-fold lower) than for wild-type EF-Tu (**Fig. 4*C***). This difference suggests binding-induced changes in the EF-Tu N-terminal peptide or EftM structure occur upon protein-protein interaction even though the EF-Tu N-terminal peptide does not appear to directly contribute to the binding affinity of the complex.

**Table 1.**
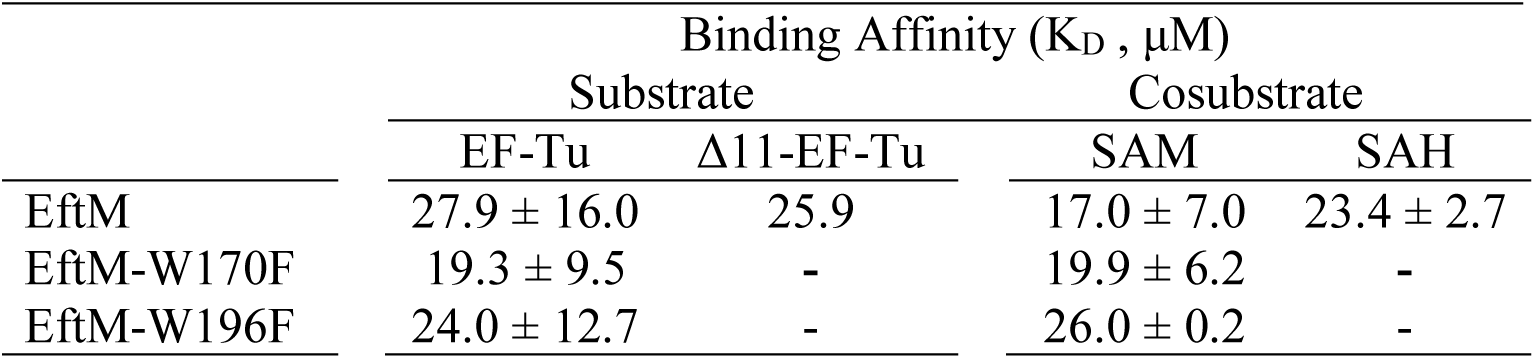
EF-Tu and SAM/ SAH binding affinities of wild-type and tryptophan variant EftM^HM4^.

| | Binding Affinity ($K_D$, μM) | | | |
| --- | --- | --- | --- | --- |
|  | Substrate |  | Cosubstrate |  |
| | EF-Tu | Δ11-EF-Tu | SAM | SAH |
| EftM | $27.9 \pm 16.0$ | 25.9 | $17.0 \pm 7.0$ | $23.4 \pm 2.7$ |
| EftM-W170F | $19.3 \pm 9.5$ | - | $19.9 \pm 6.2$ | - |
| EftM-W196F | $24.0 \pm 12.7$ | - | $26.0 \pm 0.2$ | - |

In a complementary approach, we used mass spectroscopy (MS) to discern whether peptides corresponding to the isolated N-terminal peptide sequence of *P. aeruginosa* EF-Tu could be methylated by EftM. Samples containing 10-mer peptide (^1^MAKEKFERNK^10^) alone or after incubation with EftM/ SAM were subject to matrix-assisted laser desorption/ ionization-time of flight MS analysis. In both spectra, peaks corresponding the unmethylated peptide (m/z 1280) were identified but no methylated peptides were detected (predicted m/z 1294, 1308, 1322 for mono-, di- and trimethylated peptide, respectively), even with the additional of enzyme and SAM (**Fig. 4*D***). We additionally tested a shorter peptide (^1^MAKEKF^6^) that corresponds more precisely to the sequence we predict is accommodated within the EftM peptide binding channel (see below). Again, only unmethylated peptide was detected by MS, even in the sample containing EftM and SAM (**Fig. S1*A***). Further, synthetic trimethylated peptide (^1^MAKE**K**^**Me3**^F^6^) could be detected but, as expected, was not further modified by the addition of EftM and SAM (**Fig. S1*B***).

Together, the gel filtration chromatography and ITC data demonstrating the ability of EftM to bind EF-Tu lacking its N-terminal sequence and MS evidence that EftM cannot methylate an isolated EF-Tu N-terminal peptide sequence point to a critical role for an extended surface of the globular body of EF-Tu in substrate recognition by EftM.

### EftM interacts with an extensive surface of EF-Tu

We next generated a computational model of the EftM:EF-Tu complex using the protein-protein docking software HEX 6.0 (13). To generate this model, a structure corresponding to Δ11-EF-Tu was first created by deletion of N-terminal residues in the structure of *P. aeruginosa* EF-Tu (PDB 4ZV4). Potential models of Δ11-EF-Tu bound to EftM^HM4^ generated by HEX 6.0 were ranked according to calculated energies for shape and electrostatic complementarity. The top scoring prediction was manually adjusted, e.g. by selection of optimal side chain rotamers, to remove obvious clashes before energy minimization to generate the final model of the EftM:Δ11-EF-Tu complex (**Fig. 5*A,B***).

**Fig. 5.**
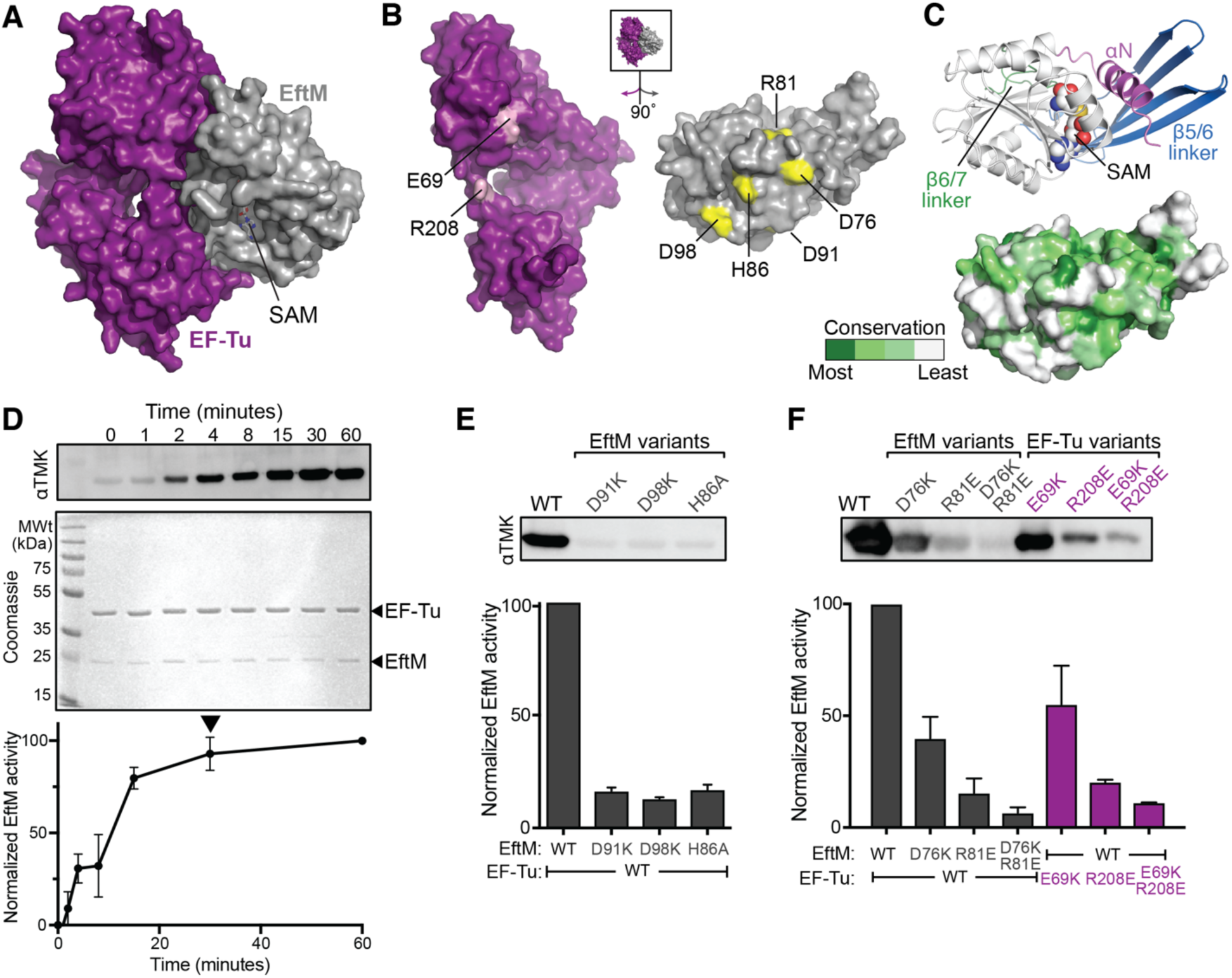
EftM interacts with an extensive surface of EF-Tu. ***A***, Docking-based model of EftM^HM4^ homology model (grey) with Δ11-EF-Tu (purple), ***B***, Residues of EftM (yellow) and EF-Tu (light purple) which were mutated to test the predicted protein-protein interface. ***C***, *Top*, a cartoon of EftM^HM4^ highlighting key structural features, colored as in **Fig. 2**. *Bottom*, surface representation with amino acid conservation of residues on the predicted EF-Tu interacting surface of EftM shown with a gradient of green-white coloring (most conserved to most variable). ***D***, Representative reaction time-course of EF-Tu methylation assay immunoblot (*top*) and Coomassie staining of methylation reactions (*center*). *Bottom*, quantification of immunoblot data from two replicate experiments. The reaction approaches completion at ∼30 min (black arrowhead) for EftM^HM4^, EF-Tu and SAM concentrations of 0.25 μM, 10 μM and 1 mM, respectively. ***E***, Representative trimethyllysine immunoblot (*top*) and quantification of replicates (*bottom*) for EF-Tu methylation with EftM^HM4^ amino acid substitutions on the interacting surface: D91K, D98K, and H86A. **F.** As *panel E* but for EftM^HM4^ variants D76K, R81E and D76K/R81E, and EF-Tu variants E69K, R208E and E69K/R208E.

The final model was assessed based on amino acid evolutionary conservation within the EftM family, revealing a moderately conserved surface of EftM^HM4^ that is predicted to interact with EF-Tu (**Fig. 5*B,C***). In contrast, the predicted solvent exposed surface of EftM exhibited lower overall conservation, whereas the predicted openings to the EF-Tu N-terminal peptide binding channel have much higher amino acid conservation as would be expected for this functionally critical region of the enzyme (**Fig. S2**). The putative EftM:Δ11-EF-Tu binding interface is stabilized by multiple electrostatic interactions made by EftM^HM4^ residues Asp76, Arg81, Asp91, and Asp98. To experimentally test the role of these residues in EF-Tu binding, each was altered with a charge reversal substitution (*i.e.* Arg to Glu or Asp to Lys). Additionally, His86 within the same surface was substituted with Ala. This group of residues is distributed across most of the predicted interaction interface (**Fig. 5*B***). Variant EftM^HM4^ proteins were expressed and purified as for the wild-type protein and their folding assessed using thermal denaturation monitored by intrinsic fluorescence. Each protein had a similar unfolding profile and inflection point melting temperature (T_i_; within ±5 °C) compared to wild-type EftM^HM4^ consistent with their correct folding. Additionally, since each substituted residue is located on the EftM^HM4^ protein surface and distant from both the SAM binding pocket and EF-Tu N-terminal peptide binding channel, we therefore ascribe any observed decrease in trimethylation activity to weakening of the binding interaction between EftM^HM4^ and its EF-Tu substrate at this predicted surface.

To measure trimethylation of EF-Tu by the variant EftM^HM4^ proteins, we used the established immunoblot assay with an anti-trimethyllysine antibody (αTMK). First, to determine the optimal reaction conditions for comparison with wild-type EftM^HM4^, a time-course activity assay was performed using wild-type EftM^HM4^ (0.25 μM) in the presence of excess EF-Tu substrate (10 μM) and SAM cosubstrate (1 mM). This experiment revealed that under the conditions used, EF-Tu trimethylation neared completion at ∼30 min (**Fig. 5*D***) and this single time point was selected for use in all subsequent methylation assays. D91K, D98K, and H86A substitutions in EftM^HM4^ resulted in a severely reduced ability to trimethylate EF-Tu (**Fig. 5*E***), with activity decreased to ∼16%, 13%, and 17%, respectively, compared to wild-type EftM^HM4^ (**Fig. 5*E***). D76K and R81E substitutions also caused a reduced ability to methylate EF-Tu, to ∼40%, and 15%, respectively, while double substitution of these residues in EftM^HM4^-D76K/R81E further lowered activity to ∼7%.

We next made individual substitutions of residues Glu69 and Arg208 in EF-Tu, which are predicted by the model of the complex to be near EftM^HM4^ residues Asp76 and Arg81, respectively. Consistent with the effects of the corresponding changes in EftM^HM4^, charge reversal substitutions E69K and R208E in EF-Tu also resulted in reduced methylation activity by wild-type EftM, dropping to ∼55% and 20%, respectively (**Fig. 5*E***). The double mutant, EF-Tu-E69K/R208E, again showed a more pronounced decrease in methylation activity (to ∼11% of wild-type), mirroring the effects of the corresponding substitutions in EftM. Collectively, the activities of the EftM^HM4^ variants with targeted substitutions, and of wild-type EftM on variant EF-Tu proteins with corresponding changes, experimentally validate the predicted protein-protein interaction interface in our computational model of the EftM:Δ11-EF-Tu complex.

### EF-Tu N-terminus is accommodated in a conserved substrate-binding channel within EftM

Using our model of the EftM:EF-Tu complex, we next asked how the EF-Tu N-terminal peptide might be positioned in EftM’s substrate binding channel. The EftM^HM4^ homology model has its substrate binding channel in a similar location to that observed in the DesVI structure but with an overall larger volume and distinct surface shape and charge distribution. The residues lining the substrate binding channel, Tyr7, Asp15, Met74, Trp170, and Trp196 (**Fig. 6*A*** and **Fig. S2*B***), are highly conserved (>90%), while Asp20 near the opening of the channel is also conserved (82.4%) in EftM family proteins. Additionally, although these residues are conserved in the EftM family, they are less highly conserved in DesVI and other paralogs, highlighting their potential role in interactions with the EF-Tu N-terminal peptide which orient Lys 5 for trimethylation. Near the center of the channel, EftM also accommodates the SAM cosubstrate in an orientation consistent with other Rossman fold methyltransferases (including DesVI), in which methyl group is oriented towards the opening of substrate channel adjacent to the modeled EF-Tu N-terminus (**Fig. 6*A***).

**Fig. 6.**
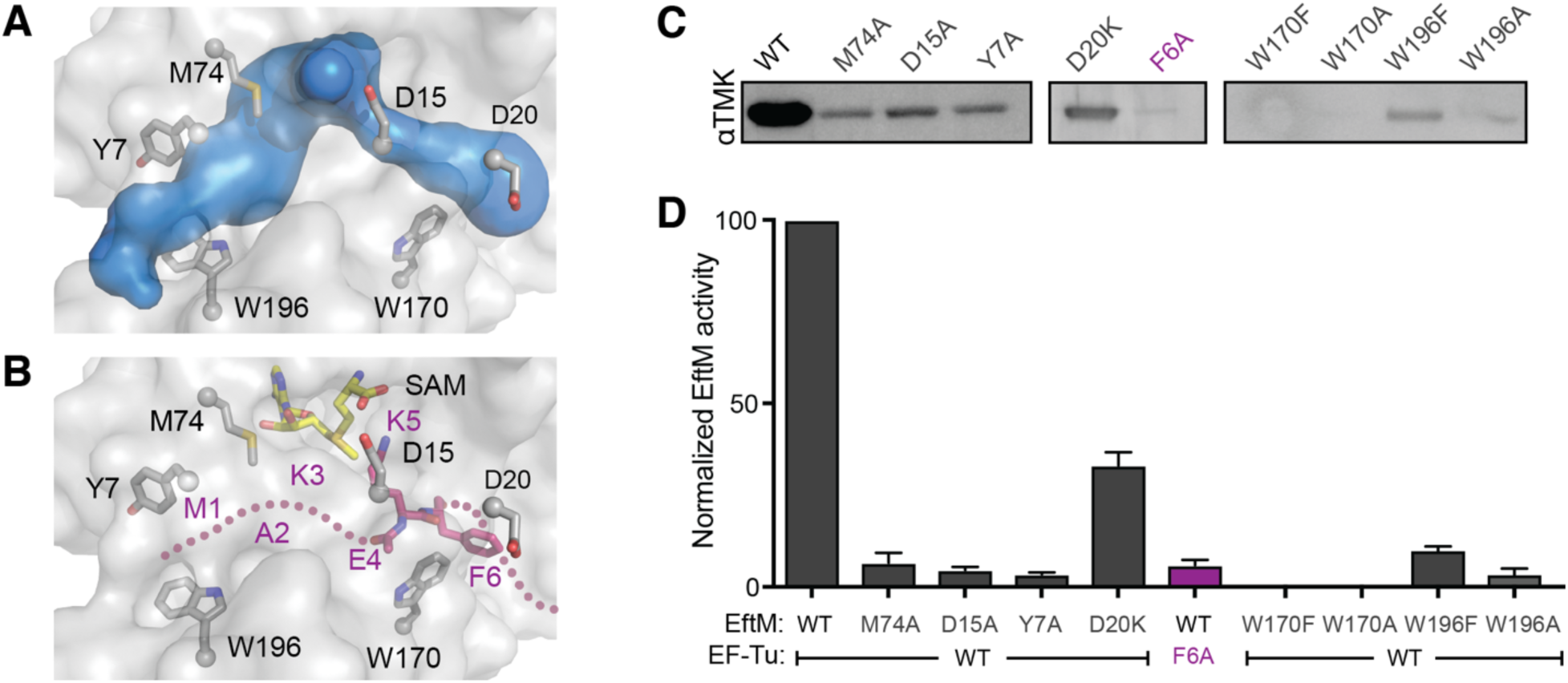
The N-terminal sequence of EF-Tu (MAKE<u>K</u>F) is stabilized by conserved residues in EftM’s substrate binding channel. ***A***, The substrate binding channel of EftM^HM4^ based on solvent space calculations using the CAVER plugin of PyMOL. EftM^HM4^ residues Tyr7, Asp15, Asp20, Met74, Trp170 and Trp196 line the substrate tunnel. ***B***, Same view but showing SAM (yellow) and three modeled EF-Tu residues Glu4 (modeled as Ala), Lys5 and Phe6 within the cavity. The approximate path of the additional N-terminal peptide backbone is marked with a dotted line. ***C***, Representative trimethyllysine immunoblots of methylation reaction with the indicated EftM^HM4^ variants and F6A substitution in EF-Tu and ***D***, quantification of immunoblot data from two replicate experiments.

The cavity within EftM that forms the EF-Tu N-terminal peptide binding channel appears large enough to accommodate the first six residues of EF-Tu (^1^MAKEKF^6^) with some distinctive surface pockets that can fit specific EF-Tu residues. We therefore next attempted to model this N-terminal sequence backbone path through the channel, positioning specific residues where possible (**Fig. 6*B***). Two EF-Tu residues, Lys5 and Phe6, could be readily positioned with minimal clashes; in other cases, while the backbone path is clear, expansion of the channel would be necessary to accommodate the residue side chains. As discussed further below, we propose that interaction of EftM and EF-Tu may induce such widening of the peptide binding channel.

To test the predicted roles of EftM residues within the substrate binding channel in recognizing and stabilizing the EF-Tu N-terminal peptide, further EftM variants were generated. Tyr7 and Met74 are located next to the predicted path of the first three residues (^1^MAK^3^) of EF-Tu may orient and stabilize the N-terminal sequence. Consistent with such a role, individual alanine substitution of each residue reduced EftM activity to 6.5% and 3.5%, respectively (**Fig. 6*C,D***). EF-Tu Lys5 is placed in the model near the EftM catalytic center, as required for its modification, and proximal to Asp15. Substitution of this EftM residue with alanine (EftM-D15A) also dramatically reduced EF-Tu methylation activity to ∼5% of the wild-type enzyme (**Fig. 6*C,D***). The modeling also places EftM Trp170 adjacent to EF-Tu Phe6 and Trp196 at the opposite end of the peptide binding channel adjacent to the EF-Tu N-terminus; as described in the following section, these residues also appear to play particularly critical roles in EftM activity. Finally, Asp20 is located just outside the peptide binding channel and could potentially interact with EF-Tu residues following Phe6. Substitution of Asp20 with alanine led to a more modest decrease in activity compared (∼40% of wild-type EftM activity; **Fig. 6*C,D***). These results suggest that interactions made to the first six N-terminal residues of EF-Tu are the most critical for positioning Lys5 for methylation.

### EftM Trp170 and Trp196 flank the EF-Tu N-terminal peptide binding channel and are critical for EftM activity

The highly conserved EftM residues Trp170 and Trp196 are positioned at each end of the EF-Tu N-terminal peptide binding channel suggesting they may perform “gatekeeping” functions at the channel openings. We therefore investigated their potential roles in cosubstrate binding and EF-Tu recognition and modification. EftM^HM4^ Trp170 and Trp196 were individually substituted to phenylalanine or alanine and the activity of each variant assessed as before. First, in the immunoblot methylation assay, both Trp170 substitutions, W170A and W170F, completely ablated EftM^HM4^ activity, while the equivalent changes at Trp196 (W196A and W196F) resulted in a very low level of trimethylation activity, ∼4% and 10%, respectively, compared to wild-type EftM^HM4^ **(Fig. 6*C***,***D*).**

We next used ITC to establish whether substitution of either residue disrupts SAM binding. The binding affinities (K_D_) of SAM and *S*-adenosyl-homocysteine (SAH) (**Table 1**) for wild-type EftM^HM4^ were first determined and found to be essentially identical to those previously measured for EftM^PAO1^ (8). Similarly, no major difference was observed in SAM binding affinity to either tryptophan to phenylalanine variant (**Table 1**), indicating that neither Trp170 nor Trp196 contribute meaningfully to cosubstrate binding.

To address whether either Trp170 or Trp196 contributes to EF-Tu binding affinity, the interaction of each tryptophan to phenylalanine variant with EftM^HM4^ was also assessed by ITC. As discussed above, wild-type EftM^HM4^ binds EF-Tu with a K_D_ of ∼28 μM and the binding affinities of both W170F and W196F are essentially identical (**Table 1**). However, as observed for EftM^HM4^ interaction with the Δ11-EF-Tu variant, a large decrease (>10-fold) in ΔH was again apparent for EftM-W170F and EftM-W196F interaction with wild-type EF-Tu (**Fig. S3**). These findings are also consistent with our earlier interpretation that a large enthalpic component of binding arises from binding-induced re-organization of the EF-Tu N-terminus into the substrate channel and, further, that these two EftM Trp residues play an important role in this process.

Toegther, these data reveal that Trp170 and Trp196 play functionally critical roles in EftM activity, with alteration of the former residue, in particular, fully ablating EF-Tu methylation. These residues do not, however, contribute directly to the binding affinity for either substrate or cosubstrate. The proximity of Trp196 to a conserved opening on the EftM surface (**Fig. S2*B***) suggests this residue could be required for interactions or conformational changes necessary for exchange of SAM/SAH during catalytic turn-over or in stabilizing the most N-terminal residues of EF-Tu, together with Tyr7 and Met74. The apparent absolute requirement for Trp170 in EftM activity together with our modeling of residues in the peptide binding channel suggest that Trp170 interacts with the highly conserved EF-Tu residue Phe6 which immediately follows the Lys5 trimethylation site to precisely orient the target residue for modification. Consistent with this idea, alteration of the EF-Tu with a F6A substitution also results in a dramatic loss of methylation by EftM (**Fig. 6*C,D***).

### Binding of the EF-Tu N-terminal peptide for Lys5 methylation

We next addressed the question of how the N-terminus of EF-Tu accesses the EftM substrate-binding channel, which in the homology model is a cavity within the enzyme with openings at each end. At least two distinct potential mechanisms can be envisaged. First, following EftM:EF-Tu interaction, the EF-Tu N-terminal sequence could enter via an adjacent opening on the EftM surface (**Fig. S2*B***) and thread into the substrate channel until fully accommodated (a “threading” model). Alternatively, binding to EF-Tu could induce conformational changes in EftM that open the peptide binding channel allowing “placement” of the ^1^MAKEKF^6^ sequence directly into its final position (**Fig. S4*A***). Given that EftM can methylate an N-terminally 6*×*His-tagged EF-Tu construct with similar efficiency to EF-Tu with an authentic N-terminus (**Fig. S5**), the latter scenario appears most plausible and we sought further evidence to more fully support this placement model and eliminate the potential alternative threading model.

To gain initial insight into potential points of flexibility in EftM that could be influenced by interaction with EF-Tu, we used the El Nemo server (14) to perform a normal mode analysis (NMA) on the EftM^HM4^ homology model structure. NMA provides a coarse model of large-scale protein movements and can identify potential hinge points and the directionality of domain movements. The NMA suggests that EftM^HM4^ can undergo a large-scale conformational change between the methyltransferase core fold and the β5/6 linker of the auxiliary domain (**Fig. 7*A*** and **Fig. S4*B***). This motion could potentially generate a more open as well as a more closed conformation of the peptide binding channel in EftM (**Fig. 7*A***), compared to the starting homology model structure. We speculate that this movement towards the “open” conformation could be induced by EF-Tu binding and thus expand the substrate-binding channel to allow direct accommodation of the ^1^MAKE<u>**K**</u>F^6^ peptide sequence.

**Fig. 7.**
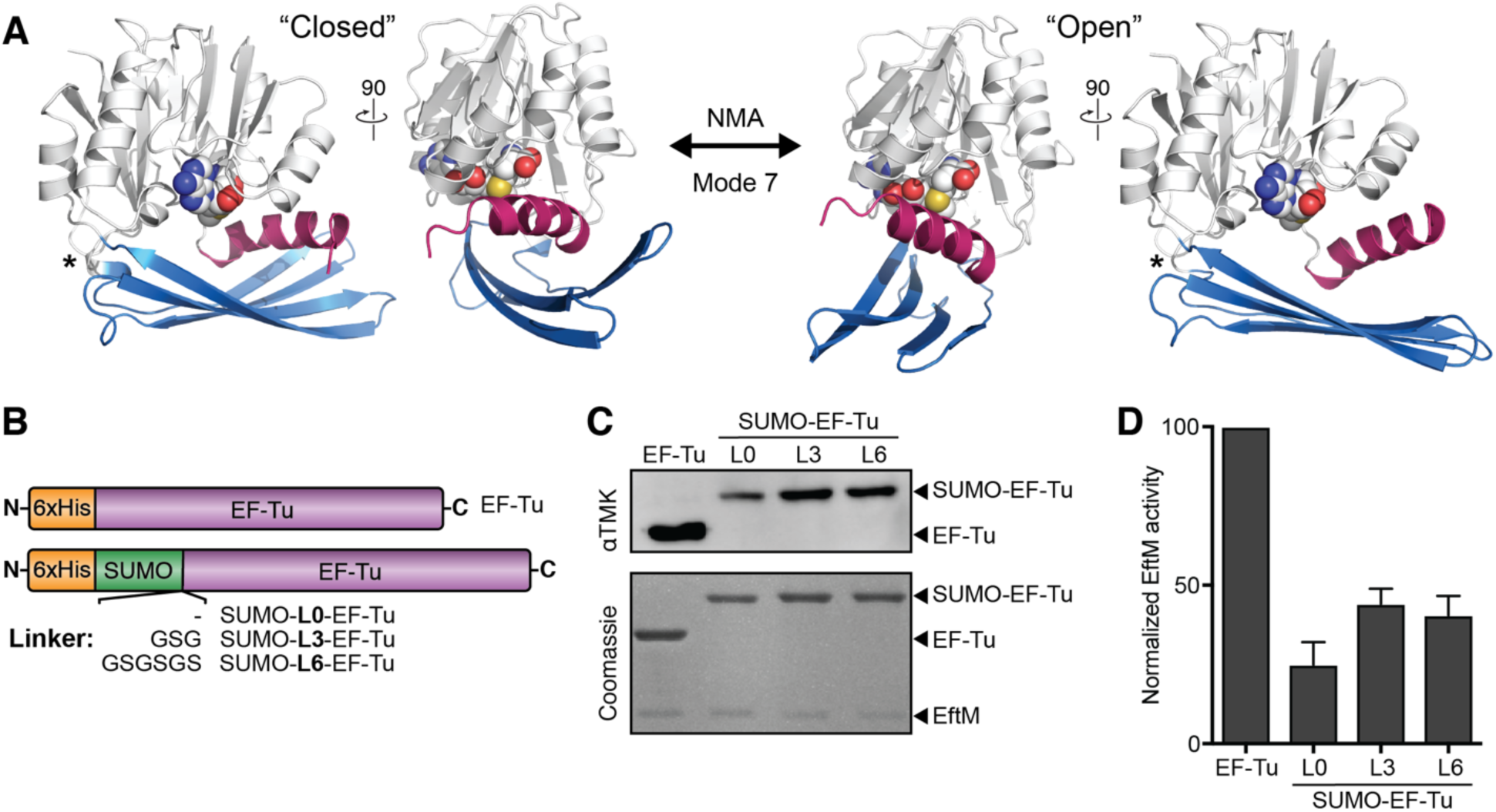
EftM flexibility allows placement of the EF-Tu N-terminal sequence into the peptide binding channel. **A**, Cartoons of two orthogonal views of the EftM homology model structure shown at the extremes of domain motions revealed by NMA Mode 7. NMA reveals that EftM may adopt structures with a more “closed” (*left*) or more “open” (*right*) form of the peptide binding channel compared to the initial homology model structure. This motion is due primarily to a hinge between the *β*5/6 linker and methyltransferase core (marked *). ***B***, Design of constructs expressing EF-Tu with an N-terminal SUMO domain fusion. ***C***, Representative trimethyllysine immunoblot of methylation reactions with wild-type EftM^HM4^ and EF-Tu and its SUMO-fusion constructs, and ***D***, quantification of two replicate assays with these proteins.

To further experimentally test the placement model, we used a construct of EF-Tu fused at its N-terminus to the SUMO domain (**Fig. 7*B***, SUMO-**L0**-EF-Tu). Our expectation was that if the channel is rigid and does not open upon EF-Tu binding then the relatively large, folded SUMO domain attached to the N-terminus of EF-Tu would preclude entry through the opening on the EftM surface (**Fig. S4*C***). Although methylation was reduced, EftM^HM4^ is still able to trimethylate Lys5 in SUMO-L0-EF-Tu with reasonable efficiency (∼30% of wild-type EftM activity; **Fig. 7*C,D***). We next increased the linker length between SUMO and EF-Tu domains, to adjust the distance between the SUMO domain and the EftM:EF-Tu complex. Two constructs with either a three (SUMO-**L3**-EF-Tu) or six (SUMO-L6-EF-Tu) residue glycine/ serine-rich linker both showed a further modest recovery of activity compared to SUMO-L0-EF-Tu (∼50% compared to wild-type EF-Tu; **Fig. 7*C***,***D***). Collectively, these results support of the model of placement of the EF-Tu N-terminus into a widened peptide binding channel in EftM upon its binding to EF-Tu.

## DISCUSSION

Some bacteria that infect the host respiratory tract present the phosphorylcholine (or, choline phosphate; ChoP) modification on their cell surface to enhance airway epithelial cell adhesion and thus promote the establishment of infection (11,15,16). *P. aeruginosa*, in contrast, does not present ChoP but instead uses the surface-exposed protein synthesis factor EF-Tu with trimethylated Lys5 as a molecular mimic of this PTM that can similarly bind to the epithelial cell platelet-activating factor receptor as a key adhesin in infection (9,11). *P. aeruginosa* EftM is the methyltransferase that trimethylates EF-Tu Lys5 in a temperature-regulated manner, corresponding to the transition from environmental to host temperature (9). Prior to the current work, the mechanism of EF-Tu substrate recognition by EftM was unclear. More broadly, the mechanisms of action bacterial lysine methyltransferases are currently not well defined in sharp contrast to their eukaryotic counterparts, which play a prominent role in epigenetics via histone protein modification (17).

Here, we showed that EftM from the chronic infection strain PAHM4 possesses conserved activity and retained thermoregulation, albeit with modestly (∼5 °C) increased thermostability, compared to the previously studied EftM^PAO1^ enzyme, making it suitable for detailed structural and mechanistic studies of this enzyme family. Our studies here revealed that the isolated EF-Tu N-terminal peptide sequence is not a suitable substrate for EftM, but instead specific substrate recognition and modification are reliant on interaction via an extended protein-protein binding interface. Comparative modeling, protein-protein docking and sequence conservation analyses were used to develop models for both the EftM^HM4^:EF-Tu interaction and accommodation of the EF-Tu N-terminal peptide (^1^MAKEKF^6^) within a binding channel in EftM. Both aspects of the modeling were tested and validated by functional analyses of EftM^HM4^ or EF-Tu variants with targeted substitutions. Our analysis also highlighted the high residue conservation of the substrate recognition channel, where the EF-Tu N-terminal peptide must be positioned for Lys5 modification. Overall, our results evoke a model in which protein-protein docking via an extended surface is required for interaction of enzyme and substrate, and subsequence modification of EF-Tu Lys5 (**Fig. 8**). Further, this interaction may induce conformational changes that open EftM peptide binding channel to allow access to the EF-Tu N-terminal peptide sequence. Within this channel several highly conserved residues, including two functionally critical Trp residues, act in concert to organize the ^1^MAKEKF^6^ sequence and thus precisely orient the Lys5 target residue for modification (**Fig. 8**).

**Fig. 8.**
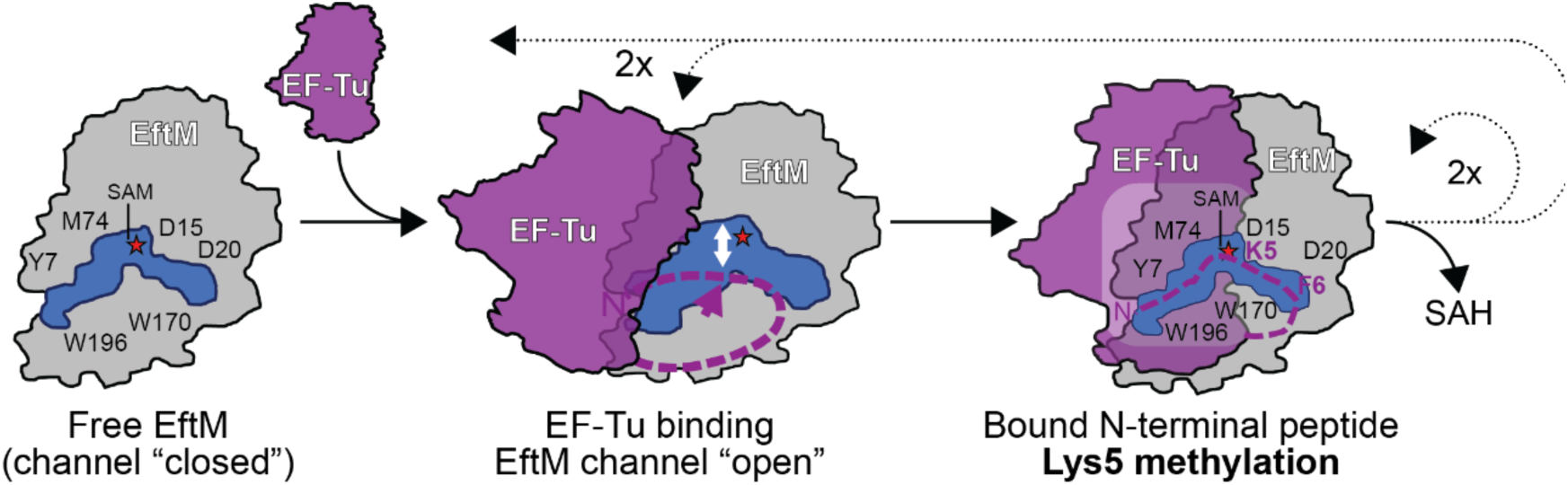
Model of EF-Tu recognition by EftM. EftM binds EF-Tu via an extended protein-protein interface, promoting conformational changes in EftM that open the peptide binding channel of the free form to a more open state that allows direct placement of the EF-Tu N-terminal peptide. Within the substrate binding channel, the N-terminal peptide of EF-Tu is oriented by a group of highly conserved residues that position Lys5 for trimethylation. The dotted arrows denoted the subsequent steps needed for additional methylation of Lys5 to generate dimethyllysine and then the final trimethyl modification. Whether SAM/SAH are recycled while EF-Tu remains fully bound (in a processive mechanism) or the substrate is partially or fully released (in a distributive mechanism) remains to be clearly defined.

The mechanism of EF-Tu recognition and Lys5 methylation by EftM revealed by our work may also have broader implications for our understanding of other lysine methyltransferases belonging to both the Class I Rossman fold- and Class II SET domain-containing methyltransferases. Specifically, our findings suggest that EftM may exploit features of molecular mechanisms common to both the Rossman fold methyltransferases, in its requirement for an extended protein surface for substrate recognition, and the SET domain methyltransferases, with its substrate binding channel that orients the modified residue within in a flexible loop or disordered protein region (18).

Despite recent discoveries of extensive methylation in bacterial proteins, including lysine methylations, and their roles in bacterial physiology and infection (4,7,19,20), there still exists a significant gap in our mechanistic understanding of bacterial lysine methyltransferases. Among bacterial lysine methyltransferases, the majority of class I Rossman fold methyltransferases recognize the globular body of their protein substrate and possess a shallow and relatively surface exposed catalytic center. For example, the ribosomal protein L11 methyltransferase, PrmA, modifies three target sites in its substrate, including two lysine residues (Lys3 and Lys39), and exploits recognition of the globular L11 protein fold and accommodates its structurally distinct target sites in a shallow catalytic center (20). This enzyme-substrate recognition mechanism thus allows the necessary flexibility in target selection for both Lys3, which resides in an unstructured N-terminal region, and Lys39, which is located within an *α*-helix. The OmpB methyltransferase in rickettsia also recognizes the globular body of its substrate to allow catalysis of multiple lysine methylations within its shallow catalytic site (21). A similar recognition mechanism is also observed in eukaryotic Class I Rossman fold methyltransferases such as DOT1L which methylates histone H3 Lys79 within the nucleosome using its N-terminal domain to bind the substrate (22). The recognition mechanism and subsequent catalysis of these methyltransferases follows a “catch and catalyze” strategy, which is mostly dependent on globular body recognition, associated conformational changes and orienting substrate in the shallow catalytic site for methylation (22). While our studies have shown that EftM also requires an extensive protein-protein interface to bind EF-Tu, its catalytic center is located within a deep substrate binding channel. This may reflect the fact that, unlike the bacterial lysine methyltransferases noted above, EftM is very specific with a single trimethylation target site at Lys5. As such, in this latter aspect of substrate recognition, EftM functions in a manner more similar to the SET domain methyltransferases.

Many SET domain methyltransferases can bind and catalyze modification of short, isolated peptide sequences containing their target site residue. The histone-lysine N-methyltransferase SETD2 specifically trimethylates Lys36 of histone H3 (H3-K36^me3^) and the enzyme is active on a 14 residue dimethylated (H3-K36^me2^) peptide as substrate (23). Upon binding to the H3 peptide, the catalytic tunnel is proposed to undergo stepwise conformational changes consistent with a placement model, accommodating and orienting the peptide (23). Similarly, EHMT1 which specifically mono- and dimethylates Lys9 of histone H3 (H3K9^me1/me2^) can recognize and methylate an 11-residue peptide from its target as the substrate, again with a mechanism consistent with the placement model (24). Other SET domain methyltransferases have additional domains fused to catalytic SET domain or act as a part of multi-protein complexes, in which both recognition of globular domain of protein(s), induced conformational rearrangements and placement of substrate motif into deep channels are observed (25). Our findings reveal that EftM has a deep peptide binding channel like the SET domain methyltransferases, and the EF-Tu N-terminal sequence is accommodated directly, likely with the aid of conformational changes induced by protein-protein interaction (**Fig. 8**).

Our current studies have revealed important details of substrate recognition by EftM and suggest that its mechanism of substrate recognition and modification is more complex than other known bacterial lysine methyltransferases. Specifically, EftM appears to more closely resemble the SET domain methyltransferases of eukaryotic origin in key aspects such as its EF-Tu N-terminal peptide binding channel. We also note that the likely catalytic center of EftM is comprised of multiple tyrosine and histidine residues, which are also found in the SET domain methyltransferases. Further investigation using high-resolution structural and careful biochemical analyses will be required to fully define the molecular basis of specific substrate as well as other aspects of EftM’s mechanism of catalysis (**Fig. 8**), such as whether EftM acts in a processive or distributive manner, as suggested by previous preliminary analysis of EftM^PAO1^ (8,9).

## EXPERIMENTAL PROCEDURES

### Computational modeling of EftM structure and the EftM:EF-Tu complex

A hybrid molecular modeling strategy was used to generate homology models of the EftM^PAO1^ and EftM^HM4^ protein structures. First, homology modeling was performed using Swiss Model (26) to generate complete structures for each protein. Inspection of these structures revealed likely incorrect modeling of the β5/6 linker region (e.g. multiple aromatic residues exposed to solvent) and this region was thus remodeled using protein threading with ITASSER (27). The model for each EftM protein was further examined in Coot (28) to confirm favorable side-chain conformations and minimize clashes, before a round of energy minimization in UCSF Chimera (29).

A structure corresponding to Δ11-EF-Tu from *P. aeruginosa* PAO1 was generated by deletion of the residues 8-11 from PDB 4ZV4 (note, the first 7 residues are disordered and were already absent in the deposited structure). This Δ11-EF-Tu structure and the EftM^HM4^ homology model were used as target and ligand, respectively, in the protein-protein docking software HEX 6.0 (13) to identify an unbiased docking interface. The models were ranked based on their total energy score, which combines calculated energies derived from both from shape complementarity and electrostatics. The top scoring model of the complex (E = 212.6 kcal/mol) was energy minimized in the UCSF Chimera (29) and then examined in Coot (28) to confirm favorable side-chain conformations. This model exhibits high electrostatic and surface shape complementarity between EftM^HM4^ and Δ11-EF-Tu, and the close placement of the EF-Tu N-terminal to EftM gives further confidence in its likely accuracy.

The CAVER 3.0.1 plugin (30) in PyMOL with shell depth and radius of 4Å each, clustering threshold of 3.5 Å, and the channel origin set at the SAM binding pocket, was used to visualize substrate binding channel in EftM.

NMA, a fast and simple method to calculate vibrational modes and protein flexibility, was performed on the EftM^HM4^ homology model using the El Nemo server (14). NMA is an implementation of an elastic network model in which the protein residues are modeled as point masses or pseudo-atoms (Cα only) connected by springs, which represent the interatomic force fields. The springs connecting each node to all other neighboring nodes are of equal strength, and only the atom pairs within a cutoff distance (within 8 Å to each other, its interaction sphere) are considered connected. The normal modes of this “ball and spring” system are calculated using a Hessian Matrix. Although this treatment is inherently simple and does not capture detailed protein dynamics, comparisons of low-frequency normal modes and the directions of large-amplitude fluctuations in molecular dynamics simulations have revealed qualitatively similar results (31). The first six NMA modes are rotational and translational, while modes 7 to 12 reflect overall long-range motions within the protein. For EftM^HM4^ the resulting modes 7 to 12 were analyzed to identify potential regions of flexibility and possible conformational changes in the protein structure.

### EftM evolutionary sequence analysis

EftM sequences were retrieved by BLAST search using EftM^PAO1^ as the query in UniProt. A set of 177 unique amino acid sequences was then selected by applying a 90% sequence identity cut-off in CD-HIT (32). Calculations of site-specific conservation were made using Geneious Prime (33) for all EftM sequences and used to calculate the position-specific Shannon entropy of the EftM protein family. The normalized Shannon entropy is given by the following expression:

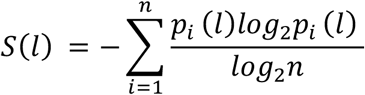

Where, *p*_*i*_*(l)* represents the frequency of *i* class of residues at position *l* in the multiple sequence alignment and n represents the number of amino acid groups depending the classification criteria. For our calculation, *n* = 20 was chosen (considering each amino acid unique). The higher entropy implies higher variability in the given position in multiple sequence alignment and *vice versa*. The calculated amino acid conservations were mapped onto the surface of the EftM^HM4^ homology model using the PDB B-factor column with a white to dark green gradient representing increasing evolutionary conservation.

### EftM and EF-Tu cloning and site-directed mutagenesis

Plasmids encoding EftM^PAO1^/ EftM^HM4^ (pColdII vector) and EF-Tu (pJP04, pDEST14-N-His_6_*tufB*) for recombinant protein expression in *Escherichia coli* or *P. aeruginosa* were previously reported (9). Site-directed mutagenesis of EftM^HM4^ and EF-Tu were performed using a mega-primer whole-plasmid PCR approach (34,35), and the variant proteins expressed from the same parent vectors as the wild-type proteins. A construct for expression of tag-free EF-Tu, was generated by PCR amplification and subcloning the *P. aeruginosa* PAO1 EF-Tu (*tufB*) coding sequence from plasmid pJP04 (9) into the pE-SUMO vector (LifeSensors). This construct produces His_6_-SUMO-EF-Tu protein (“SUMO-L0-EF-Tu”) from which the authentic EF-Tu N-terminus is generated by cleavage of the His_6_-SUMO tag by the ubiquitin-like protease (Ulp). Equivalent constructs with linker sequences extended by three or six additional amino acids between the SUMO domain and EF-Tu (“SUMO-L3-EF-Tu” and “SUMO-L6-EF-Tu”, respectively) were generated using the mega-primer whole-plasmid PCR mutagenesis approach

### EftM and EF-Tu expression and purification

Cultures of *E. coli* BL21(DE3) transformed with pColdII-EftM^HM4^ or pColdII-EftM^PAO1^ were grown at 37 °C in lysogeny broth (LB) supplemented with ampicillin (100 μg/ml) to mid-log growth (OD_600_ 0.4-0.6). After cold-shock in an ice bath for 30 minutes, protein expression was induced by addition of isopropyl-β-D-thiogalactopyranoside (IPTG; 0.5 mM) and cultures were grown for an additional 18 hours at 15 °C. Cell pellets were resuspended in 50 mM Tris, pH 7.5, lysis buffer containing 250 mM ammonium chloride, 10 mM magnesium acetate, 20 mM imidazole, 2 mM β-mercaptoethanol and 20% glycerol. Following cell lysis by sonication, soluble proteins were applied to a HisTrap-FF column (GE Healthcare) pre-equilibrated with 50 mM Tris, pH 7.5, buffer containing 150 mM sodium chloride, 5 mM magnesium chloride, 2 mM β-mercaptoethanol, 20% glycerol and 20 mM imidazole. Bound proteins were eluted using a linear gradient of imidazole (20-250 mM) in the same buffer solution. Fractions containing EftM^HM4^ or EftM^PAO1^ were pooled and the protein further purified by gel filtration chromatography on a Superdex200 10/300 column (GE Healthcare) equilibrated with gel filtration buffer containing 50 mM Tris pH 7.5, 150 mM sodium chloride, 5 mM β-mercaptoethanol and 20% glycerol.

For expression of His_6_-EF-Tu, *P. aeruginosa* PAO1Δ*eftM* cells were freshly streaked on a LB-agar plate. A single colony was used to inoculate LB (50 ml) and the culture incubated at 37 °C for ∼8 hours (to OD_600_ ∼ 0.8). Cells were then pelleted in microcentrifuge tubes and washed three times with 300 mM sucrose (1 mL) at room temperature. Sucrose-washed cells (50 μl) were transformed with pJPO04 plasmid (1.5 μl) by electroporation and allowed to recover in SOC medium (1 ml) at 37°C for one hour before plating on LB-agar plates containing 300 μg/mL carbenicillin. A single colony was then grown in LB supplemented with 0.1% arabinose and 300 μg/ml carbenicillin for 18 hours at 25 °C before harvesting. Expression of His_6_-Δ11-EF-Tu was accomplished in *E. coli* BL21-AI cells transformed with pDEST14-Δ-11tufB N-His6 fusion in ZYM-5052 medium supplemented with 0.2% arabinose (weight/volume) and carbenicillin (100 μg/mL) and incubated with shaking at 25°C for 14 hours. The His_6_-EF-Tu and His_6_-Δ11EF-Tu proteins were extracted and purified using the same chromatographic procedures as described above for the EftM proteins.

SUMO-fusion EF-Tu proteins were expressed in *E. coli* BL21 (DE3) transformed with the pE-SUMO-EF-Tu vector (L0, L3 or L6) and grown at 37 °C in LB supplemented with ampicillin (100 μg/ml) to mid-log phase (OD_600_ ∼0.4-0.6). Protein expression was induced by addition of IPTG (1 mM) and cultures were grown for 4-5 hours at 37 °C. SUMO-fusion EF-Tu proteins were purified using the same chromatographic procedures as described above for EftM. Where needed, the His_6_-SUMO tag was removed from partially purified His_6_-SUMO-L0-EF-Tu following the Ni^2+^-affinity chromatography step by overnight digestion with Ulp (0.1 μg protease per 1.25 mg fusion protein) at 37 °C. The free His_6_-SUMO tag was removed from EF-Tu by passing the protease cleavage reaction back over a HisTrap-FF column before further purification of EF-Tu by gel filtration chromatography.

### In vitro EF-Tu methylation assay

For EftM^PAO1^ and EftM^HM4^ time course methylation assay at different temperatures (25 °C and 37 °C), methylation reactions contained 10 μM of both EftM and EF-Tu and 1 mM SAM in gel filtration buffer (50 mM Tris pH 7.5, 150 mM sodium chloride, 5 mM β-mercaptoethanol and 20% glycerol). For assays investigating the thermolability of EftM^HM4^ (**Fig. 1*B***), enzyme was pre-incubated at 25, 37, or 42 °C for 0, 2 5, 10 or 20 minutes prior to performing the methylation assay. Reactions were quenched in 2X SDS-PAGE buffer and resolved on a 12% SDS-PAGE gel and immunoblotted as described below.

Methylation assays for testing the substrate recognition model were performed at 25 °C and contained EF-Tu (10 μM), EftM (0.25 μM) and 1 mM SAM in the same buffer as above. For the methylation reaction time-course with wild-type EftM^HM4^ (**Fig. 5*D***), samples were removed at each time point (2, 4, 8, 15, 30 and 60 minutes) for analysis. All subsequent EF-Tu methylation assays with wild-type and variant EftM^HM4^ and EF-Tu proteins were performed for 30 minutes. Reactions were quenched in 2*×*SDS-PAGE buffer and resolved on two 10% SDS-PAGE gel, one for immunoblotting and a second for staining with Coomassie. After transfer onto a PDVF membrane, blots were blocked using 5% non-fat milk in phosphate buffered saline with 2% Tween-20 at room temperature and then probed overnight at 4 °C with a rabbit polyclonal anti-trimethyllysine antibody (Immunchem, ICP0601) diluted 1:10000 in the same buffer. Blots were probed for 1 hour at room temperature with a horseradish peroxidase-conjugated goat anti-rabbit secondary antibody (Sigma-Aldrich; A0545) diluted 1:10000 in the same buffer. The blots were treated with enhanced chemiluminescence reagent (Thermo Fisher) and the intensity of bands was analyzed on a Bio-Rad ChemiDoc™ Imager and quantified using ImageQuant software (GE Healthcare). Two replicates for each methylation reaction were performed and quantified, and figures show one representative blot for each protein variant.

### Protein stability measurement

Differential scanning fluorimetry (DSF) was used to compare the thermal stability of EftM^PAO1^ and EftM^HM4^. Proteins samples (24 μM) were subject to a linear temperature gradient (0.5 °C/minute from 25-75 °C) and fluorescence of the hydrophobic residue-binding dye SYPRO Orange (5000-fold dilution) measured on a Step One Plus Real-Time PCR instrument (Applied Biosystems). DSF measurements were performed for each protein in the absence and presence of SAM (150 μM) in three replicates, together with corresponding control experiments without protein under each condition. The first derivative of the melting curve from each replicate was calculated using GraphPad Prism software to determine the mid-point melting temperature (T_m_). The data for EftM^PAO1^ were previously reported in (8) and shown here for comparison with EftM^HM4^.

For protein quality control and analysis of variant protein folding compared to the corresponding wild-type protein, intrinsic fluorescence (at 350 and 330 nm) was monitored during unfolding over a linear temperature gradient (35-95 °C) over 3 minutes on a Tycho NT.6 instrument (NanoTemper). The temperature at which a transition occurs in a plot of the first derivative of the fluorescence ratio (350/330 nm), termed inflection temperature (T_i_), was determined by the instrument software and compared for each protein.

### Analysis of protein binding by gel filtration chromatography

Gel filtration chromatography was performed to qualitatively assess the binding of EF-Tu or Δ11-EF-Tu with EftM^HM4^. EftM and EF-Tu were mixed in a 500 μl reaction and resolved on a Superdex 200 column (GE Healthcare) using 50 mM Tris, pH 7.5, buffer containing 150 mM sodium chloride, 5 mM β-mercaptoethanol and 20% glycerol. One ml fractions were collected and a sample of each run on an SDS-PAGE gel and stained with Coomassie to confirm coelution of EftM and EF-Tu in the first peak.

### Isothermal titration calorimetry

SAM and SAH were each dissolved in gel filtration buffer to 1 mM final concentration for EftM-cosubstrate binding experiments. EF-Tu and Δ11-EF-Tu were concentrated to ∼500 μM in the same buffer for protein-protein binding experiments. These ligands (SAM, SAH or EF-Tu proteins) were titrated into EftM (30-50

μM) in 16 × 2.4 μl injections using an Auto-iTC_200_ microcalorimeter (Malvern/MicroCal) at 25 °C. After accounting for the heat of dilution by subtraction of the residual heat measured at the end of the titration, the data were fit using a model for one set of sites to determine the binding affinity (Kd).

### Mass spectrometry analysis of EF-Tu peptide modification

Reactions containing 10 μM EftM, 10 μM peptide, and 1 mM SAM were incubated at 25 °C for 30 minutes. Peptide or methylation reaction was spotted onto the source target with 10 mg/ml α-cyano-4-hydroxycinnamic acid matrix in a 50% acetonitrile/ 0.1% trifluoracetic acid solution. MS data were collected in positive ion mode on a Bruker UltraFlex II MALDI instrument.

## Acknowledgements

We thank Dr. Dina Moustafa for assistance with protein expression. We are also grateful to other members of the Conn and Goldberg groups and those of Dr. Christine Dunham at Emory University for comments on the manuscript and helpful discussions throughout the course of this work.

## FOOTNOTES

This work was supported by the Cystic Fibrosis Foundation grant GOLDBE17P0 (to JBG and GLC) and postdoctoral fellowship DEY18F0 (to DD). SMP was supported in part by a training grant from the National Institute of Allergy and Infectious Diseases of the NIH to the Antimicrobial Resistance and Therapeutic Discovery Training Program of Emory University (T32-AI106699).

## Abbreviations

DSF: differential scanning fluorimetry
EftM: EF-Tu methyltransferase
EF-Tu: elongation factor thermo-unstable
IPTG: isopropyl-β-D-thiogalactopyranoside
ITC: isothermal titration calorimetry
LB: lysogeny broth
NMA: normal mode analysis
PTM: posttranslational modification
PAHM4: *P. aeruginosa* strain HM4
PAO1: *P. aeruginosa* strain PAO1
SAH: S-adenosyl-homocysteine
SAM: S-adenosyl-L-methionine
T_i_: inflection temperature.

## REFERENCES

1. Bastos, P. A. D., da Costa, J. P., and Vitorino, R. (2017) A glimpse into the modulation of post-translational modifications of human-colonizing bacteria. J Proteomics 152, 254–275

2. Macek, B., Forchhammer, K., Hardouin, J., Weber-Ban, E., Grangeasse, C., and Mijakovic, (2019) Protein post-translational modifications in bacteria. Nat Rev Microbiol

3. Cain, J. A., Solis, N., and Cordwell, S. J. (2014) Beyond gene expression: the impact of protein post-translational modifications in bacteria. J Proteomics 97, 265–286

4. Garcia-Fontana, C., Corral Lugo, A., and Krell, T. (2014) Specificity of the CheR2 methyltransferase in *Pseudomonas aeruginosa* is directed by a C-terminal pentapeptide in the McpB chemoreceptor. Science signaling 7, ra34

5. Sharma, G., Upadhyay, S., Srilalitha, M., Nandicoori, V. K., and Khosla, S. (2015) The interaction of mycobacterial protein Rv2966c with host chromatin is mediated through non-CpG methylation and histone H3/H4 binding. Nucleic Acids Res 43, 3922–3937

6. Yaseen, I., Choudhury, M., Sritharan, M., and Khosla, S. (2018) Histone methyltransferase SUV39H1 participates in host defense by methylating mycobacterial histone-like protein HupB. EMBO J 37, 183–200

7. Abeykoon, A., Wang, G., Chao, C. C., Chock, P. B., Gucek, M., Ching, W. M., and Yang, D. C. (2014) Multimethylation of Rickettsia OmpB catalyzed by lysine methyltransferases. J Biol Chem 289, 7691–7701

8. Owings, J. P., Kuiper, E. G., Prezioso, S. M., Meisner, J., Varga, J. J., Zelinskaya, N., Dammer, E. B., Duong, D. M., Seyfried, N. T., Alberti, S., Conn, G. L., and Goldberg, J. B. (2016) Pseudomonas aeruginosa EftM Is a Thermoregulated Methyltransferase. J Biol Chem 291, 3280–3290

9. Barbier, M., Owings, J. P., Martinez-Ramos, I., Damron, F. H., Gomila, R., Blazquez, J., Goldberg, J. B., and Alberti, S. (2013) Lysine trimethylation of EF-Tu mimics platelet-activating factor to initiate Pseudomonas aeruginosa pneumonia. MBio 4, e00207–00213

10. Prezioso, S. M., Duong, D. M., Kuiper, E. G., Deng, Q., Alberti, S., Conn, G. L., and Goldberg, J. B. (2019) Trimethylation of Elongation Factor-Tu by the Dual Thermoregulated Methyltransferase EftM Does Not Impact Its Canonical Function in Translation. Sci Rep 9, 3553

11. Barbier, M., Oliver, A., Rao, J., Hanna, S. L., Goldberg, J. B., and Alberti, S. (2008) Novel phosphorylcholine-containing protein of Pseudomonas aeruginosa chronic infection isolates interacts with airway epithelial cells. J Infect Dis 197, 465–473

12. Burgie, E. S., and Holden, H. M. (2008) Three-dimensional structure of DesVI from Streptomyces venezuelae: a sugar N,N-dimethyltransferase required for dTDP-desosamine biosynthesis. Biochemistry 47, 3982–3988

13. Macindoe, G., Mavridis, L., Venkatraman, V., Devignes, M. D., and Ritchie, D. W. (2010) HexServer: an FFT-based protein docking server powered by graphics processors. Nucleic Acids Res 38, W445–449

14. Suhre, K., and Sanejouand, Y. H. (2004) ElNemo: a normal mode web server for protein movement analysis and the generation of templates for molecular replacement. Nucleic Acids Res 32, W610–614

15. Clark, S. E., and Weiser, J. N. (2013) Microbial modulation of host immunity with the small molecule phosphorylcholine. Infect Immun 81, 392–401

16. Smani, Y., Docobo-Perez, F., Lopez-Rojas, R., Dominguez-Herrera, J., Ibanez-Martinez, J., and Pachon, J. (2012) Platelet-activating factor receptor initiates contact of Acinetobacter baumannii expressing phosphorylcholine with host cells. J Biol Chem 287, 26901–26910

17. Hyun, K., Jeon, J., Park, K., and Kim, J. (2017) Writing, erasing and reading histone lysine methylations. Exp Mol Med 49, e324

18. Del Rizzo, P. A., and Trievel, R. C. (2014) Molecular basis for substrate recognition by lysine methyltransferases and demethylases. Biochim Biophys Acta 1839, 1404–1415

19. Alvarez-Venegas, R. (2014) Bacterial SET domain proteins and their role in eukaryotic chromatin modification. Front Genet 5, 65

20. Demirci, H., Gregory, S. T., Dahlberg, A. E., and Jogl, G. (2008) Multiple-site trimethylation of ribosomal protein L11 by the PrmA methyltransferase. Structure 16, 1059–1066

21. Abeykoon, A. H., Noinaj, N., Choi, B. E., Wise, L., He, Y., Chao, C. C., Wang, G., Gucek, M., Ching, W. M., Chock, P. B., Buchanan, S. K., and Yang, D. C. (2016) Structural Insights into Substrate Recognition and Catalysis in Outer Membrane Protein B (OmpB) by Protein-lysine Methyltransferases from Rickettsia. J Biol Chem 291, 19962–19974

22. Valencia-Sanchez, M. I., De Ioannes, P., Wang, M., Vasilyev, N., Chen, R., Nudler, E., Armache, J. P., and Armache, K. J. (2019) Structural Basis of Dot1L Stimulation by Histone H2B Lysine 120 Ubiquitination. Mol Cell 74, 1010–1019 e1016

23. Yang, S., Zheng, X., Lu, C., Li, G. M., Allis, C. D., and Li, H. (2016) Molecular basis for oncohistone H3 recognition by SETD2 methyltransferase. Genes Dev 30, 1611–1616

24. Wu, H., Min, J., Lunin, V. V., Antoshenko, T., Dombrovski, L., Zeng, H., Allali-Hassani, A., Campagna-Slater, V., Vedadi, M., Arrowsmith, C. H., Plotnikov, A. N., and Schapira, M. (2010) Structural biology of human H3K9 methyltransferases. PLoS One 5, e8570

25. Kuzmichev, A., Nishioka, K., Erdjument-Bromage, H., Tempst, P., and Reinberg, D. (2002) Histone methyltransferase activity associated with a human multiprotein complex containing the Enhancer of Zeste protein. Genes Dev 16, 2893–2905

26. Waterhouse, A., Bertoni, M., Bienert, S., Studer, G., Tauriello, G., Gumienny, R., Heer, F. T., de Beer, T. A. P., Rempfer, C., Bordoli, L., Lepore, R., and Schwede, T. (2018) SWISS-MODEL: homology modelling of protein structures and complexes. Nucleic Acids Res 46, W296–W303

27. Yang, J., Yan, R., Roy, A., Xu, D., Poisson, J., and Zhang, Y. (2015) The I-TASSER Suite: protein structure and function prediction. Nat Methods 12, 7–8

28. Emsley, P., Lohkamp, B., Scott, W. G., and Cowtan, K. (2010) Features and development of Coot. Acta Crystallogr D Biol Crystallogr 66, 486–501

29. Pettersen, E. F., Goddard, T. D., Huang, C. C., Couch, G. S., Greenblatt, D. M., Meng, E. C., and Ferrin, T. E. (2004) UCSF Chimera--a visualization system for exploratory research and analysis. J Comput Chem 25, 1605–1612

30. Jurcik, A., Bednar, D., Byska, J., Marques, S. M., Furmanova, K., Daniel, L., Kokkonen, P., Brezovsky, J., Strnad, O., Stourac, J., Pavelka, A., Manak, M., Damborsky, J., and Kozlikova, B. (2018) CAVER Analyst 2.0: analysis and visualization of channels and tunnels in protein structures and molecular dynamics trajectories. Bioinformatics 34, 3586–3588

31. Alexandrov, V., Lehnert, U., Echols, N., Milburn, D., Engelman, D., and Gerstein, M. (2005) Normal modes for predicting protein motions: a comprehensive database assessment and associated Web tool. Protein Sci 14, 633–643

32. Fu, L., Niu, B., Zhu, Z., Wu, S., and Li, W. (2012) CD-HIT: accelerated for clustering the next-generation sequencing data. Bioinformatics 28, 3150–3152

33. Kearse, M., Moir, R., Wilson, A., Stones-Havas, S., Cheung, M., Sturrock, S., Buxton, S., Cooper, A., Markowitz, S., Duran, C., Thierer, T., Ashton, B., Meintjes, P., and Drummond, A. (2012) Geneious Basic: an integrated and extendable desktop software platform for the organization and analysis of sequence data. Bioinformatics 28, 1647–1649

34. Miyazaki, K. (2011) MEGAWHOP cloning: a method of creating random mutagenesis libraries via megaprimer PCR of whole plasmids. Methods Enzymol 498, 399–406

35. Savic, M., Sunita, S., Zelinskaya, N., Desai, P. M., Macmaster, R., Vinal, K., and Conn, G. L. (2015) 30S Subunit-dependent activation of the Sorangium cellulosum So ce56 aminoglycoside resistance-conferring 16S rRNA methyltransferase Kmr. Antimicrob Agents Chemother 59, 2807–2816

