## Supporting Information for "Substrate recognition by the *Pseudomonas aeruginosa* EF-Tu methyltransferase EftM"

Running title: *EF-Tu recognition by EftM*

†These authors contributed equally to this work.

---

**Supplementary Figures S1-S5**

**Fig. S1** EftM does not methylate the isolated [1]MAKEKF[6] peptide from the EF-Tu N-terminus.

**Fig. S2** Surface residue conservation in $EftM^{HM4}$.

**Fig. S3** ITC analysis of EftM Trp170 and Trp196 variant binding to EF-Tu.

**Fig. S4** NMA suggests EftM protein flexibility may facilitate direct "placement" of the EF-Tu N-terminal sequence into the peptide binding channel.

**Fig. S5** Comparison of EF-Tu vs 6×His-tagged EF-Tu as substrates for Lys5 trimethylation by EftM.

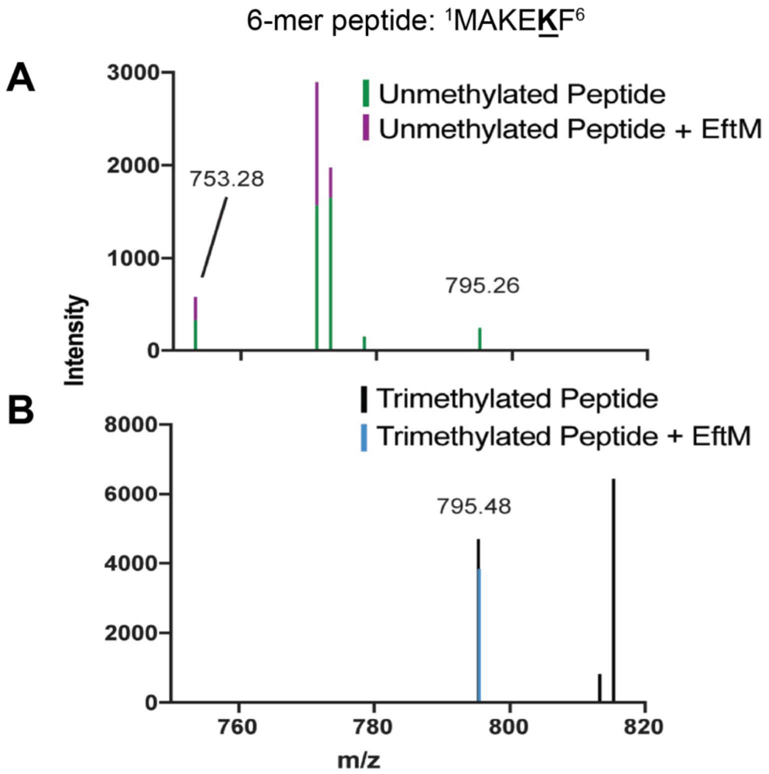


**Fig. S1 EftM does not methylate the isolated $^{1}$MAKEKF$^{6}$ peptide from the EF-Tu N-terminus.** MS analysis of the N-terminal six amino acid peptide of EF-Tu after treatment in the absence or presence of EftM/ SAM ("+EftM") using ***A***, unmodified 6-mer peptide (show top) and ***B***, Lys5 trimethylated 6-mer peptide (MAKEK$^{Me3}$F).

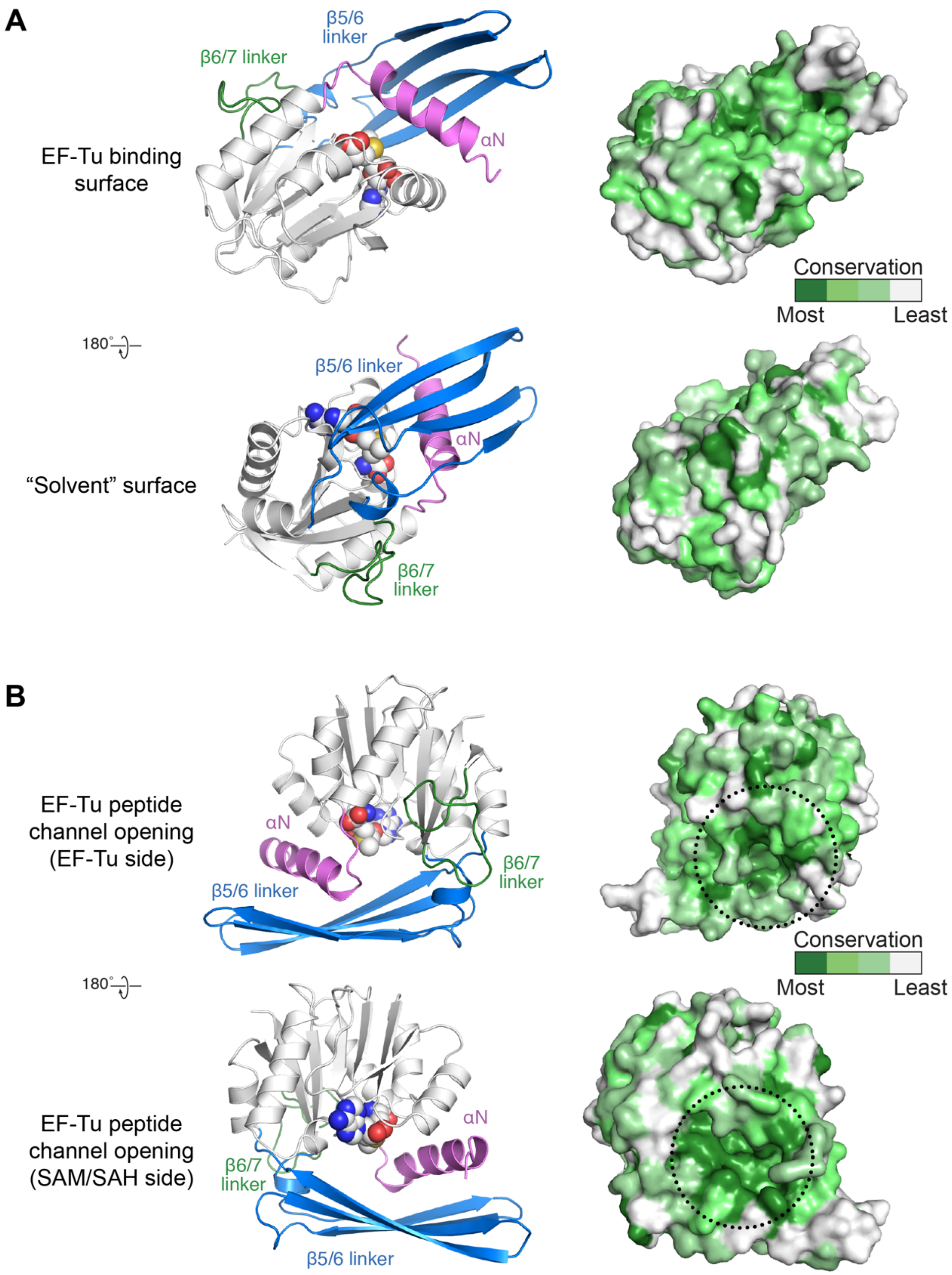


**Fig. S2 Surface conservation of EftM$^{HM4}$.** EftM$^{HM4}$ homology model structure shown as cartoon (*left*, structural features colored coded as in **Fig. 2**) and as surface representation color-coded by residue conservation (*right*). ***A***, Views of the predicted EF-Tu binding surface (*top*; same view as in **Fig. 5*C***) and the opposite, solvent exposed surface (*bottom*). ***B***, Views of the access points to the EF-Tu N-terminal peptide binding channel, proximal to the end of the EF-Tu N-terminal peptide (residue 10; *top*) and for potential cycling of SAM/ SAH (*bottom*).

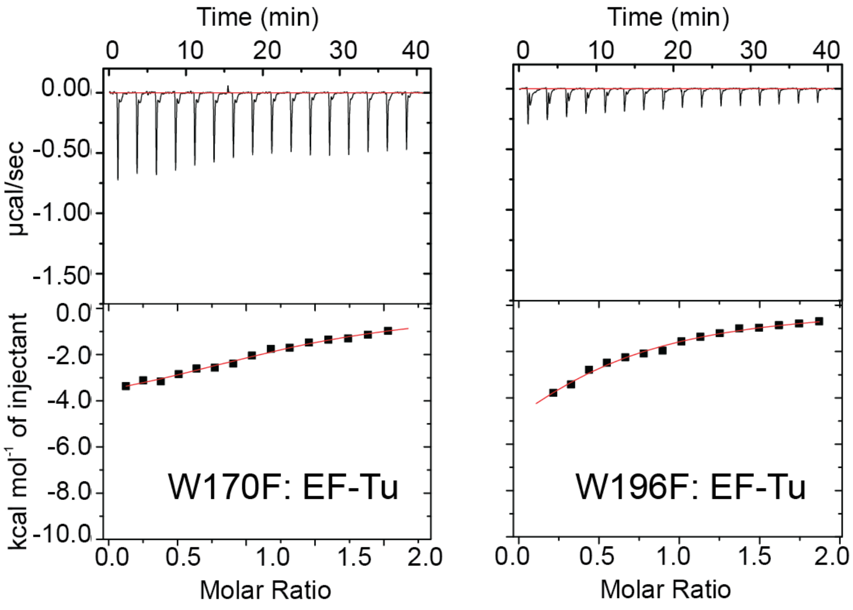


**Fig. S3 ITC analysis of EftM Trp170 and Trp196 variant binding to EF-Tu.** Example titrations of EF-Tu into EftM W170F (*left*) and W196F (*right*) variants.

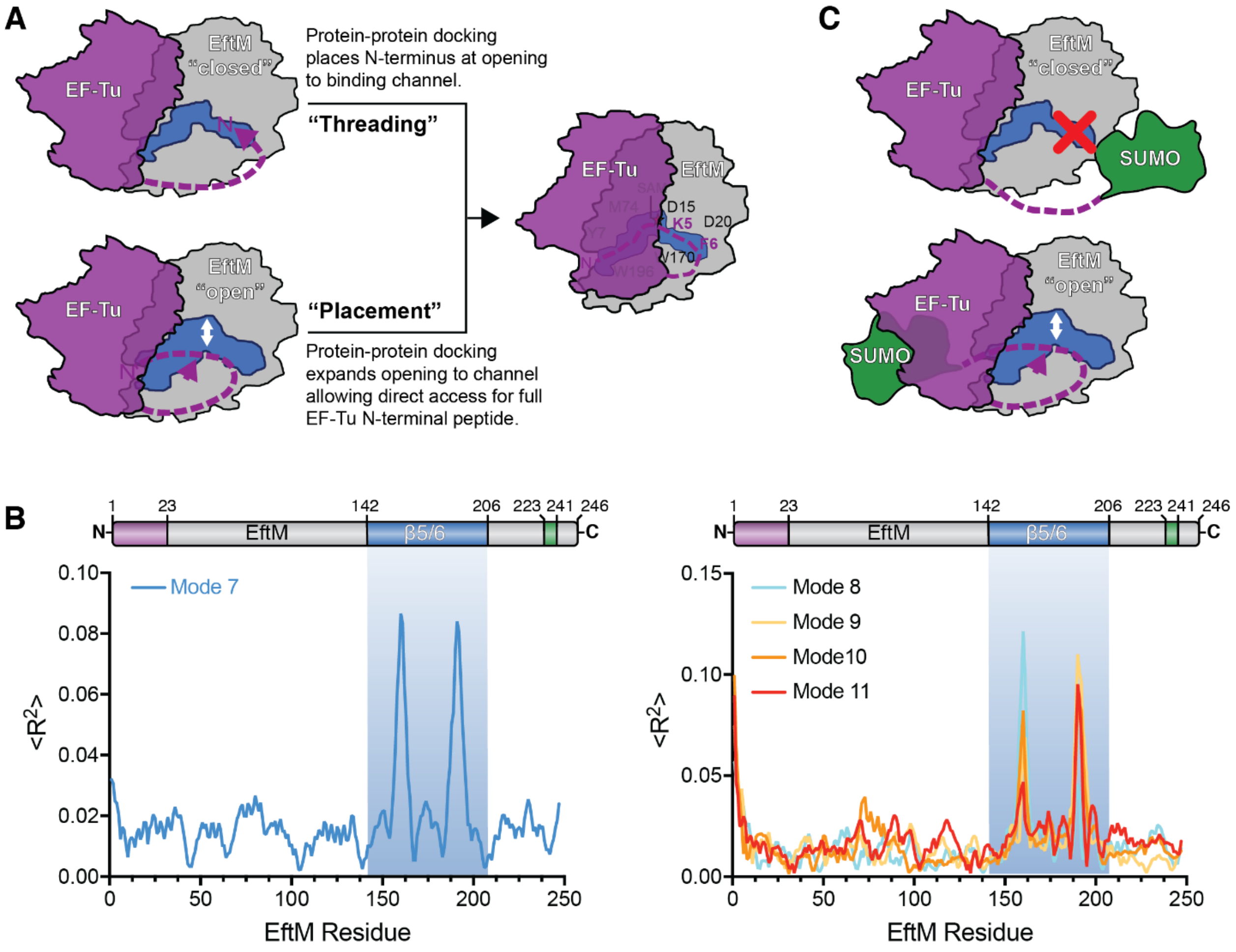


**Fig. S4 NMA suggests EftM protein flexibility may facilitate direct "placement" of the EF-Tu N-terminal sequence into the peptide binding channel**. ***A***, Potential models for how the EF-Tu N-terminal peptide ($^{1}$MAKEKF$^{6}$) accesses the EftM peptide binding channel. The white double-headed arrow indicates the need for opening of the channel to allow direct "placement" of the N-terminal sequence into its binding site. ***B***, NMA displacement plots for Mode 7 (*left*) and all other relevant modes (*right*). EftM domain boundaries are indicated by the bars above the plots, color-coded as in **Fig. 2**. ***C***, Fusion of an additional folded protein domain, such as SUMO, is predicted to ablate activity by the "threading" mechanism but should be tolerated in a "placement" mechanism.

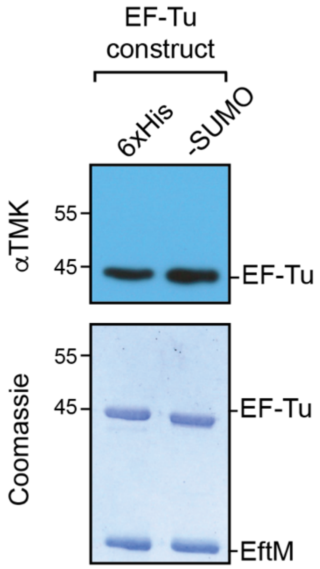


**Fig. S5 Comparison of EF-Tu vs 6×His-tagged EF-Tu as substrates for Lys5 trimethylation by EftM**. Immunoblot EftM methyltransferase assay with the N-terminally 6×His-tagged EF-Tu construct (6×His) used in most experiments in this work and a construct with an authentic N-terminus generated by cleavage SUMO-L0-EF-Tu with Ulp1.
